# Allelic variation in the *Arabidopsis* TNL CHS3/CSA1 immune receptor pair reveals two functional regulatory modes

**DOI:** 10.1101/2022.05.10.491418

**Authors:** Yu Yang, Nak Hyun Kim, Volkan Cevik, Pierre Jacob, Li Wan, Oliver J. Furzer, Jeffery L. Dangl

## Abstract

Some plant NLR immune receptors are encoded in head-to-head pairs that function together. Alleles of the NLR pair CHS3/CSA1 form three clades. The clade 1 sensor CHS3 contains an integrated domain (ID) with homology to regulatory domains, which is lacking in clades 2 and 3. We defined two regulatory modes for CHS3/CSA1 pairs. One is likely mediated by effector binding to the clade 1 ID of CHS3 and the other relies on CHS3/CSA1 pairs from all clades detecting effector modification of an associated pattern recognition receptor. We suggest that an ancestral Arabidopsis CHS3/CSA1 pair gained a second recognition specificity and regulatory mechanism through ID acquisition, while retaining its original specificity as a ‘Guard’ against perturbation of pattern recognition receptor targeting by a pathogen effector. This likely comes with a cost, since both ID and non-ID alleles of the pair persist in diverse Arabidopsis populations through balancing selection.

**Summary:** We dissect a novel case where two regulatory modes emerged across three clades of the co-evolved CHS3/CSA1 plant immune receptor pairs, which features recruitment of an integrated domain (ID) into the clade 1 CHS3 alleles. Pre- and post-ID integration alleles maintain functionality; balancing selection maintains both in the Arabidopsis pan-genome.

## Introduction

Plants evolved a two-layered immune system to defend themselves against pathogen infection (Dodds and Rathjen, 2016; Jones and Dangl, 2006; Zhou and Zhang, 2020). Extracellular microbe/pathogen-associated molecular patterns (MAMPs/PAMPs) are recognized by cell surface-localized pattern-recognition receptors (PRRs) and initiate pattern-triggered immunity (PTI) (Boller and Felix, 2009; Couto and Zipfel, 2016; Macho and Zipfel, 2014). Some effectors that are delivered into host plants by pathogens to circumvent PTI and promote pathogenesis (Mukhtar et al., 2011; Weßling et al., 2014) are recognized by intracellular nucleotide-binding (NB), leucine-rich repeat (LRR) receptors (NLRs), leading to effector-triggered immunity (ETI), often culminating in a hypersensitive cell death response (HR) (Cui et al., 2015; Jones et al., 2016; Bayless and Nishimura, 2020). The two tiers of immune signaling pathway mutually potentiate each other (Ngou et al., 2021; Pruitt et al., 2021; Tian et al., 2021; Yuan et al., 2021).

NLRs detect pathogen effectors either directly as ligands bound to canonical NLR domains or to non-canonical motifs like integrated domains (IDs). Alternatively, NLRs indirectly monitor effector-mediated modification of host target proteins, known as guardees, or decoys of guardees (Jones et al., 2016; Van der Hoorn and Kamoun, 2008). NLRs are divided into two major structural groups either carrying Toll/interleukin-1 (TIR) or coiled-coiled (CC) domains at the N terminus, termed TNL (TIR-NLR) or CNL (CC-NLR), respectively (Jones et al., 2016; Bayless and Nishimura, 2020). Upon effector activation, NLRs oligomerize and form “resistosome” complexes to activate calcium influx (CNLs) or to activate TNL NADase activity. The latter generates small molecules that activate an EDS1 anchored complex to initiate cell death and defense through the calcium channel function of a class of ‘helper’ NLRs called RNLs (Bi et al., 2021; Huang et al., 2022; Jacob et al., 2021; Martin et al., 2020; Ma et., 2020; Jubic et al., 2019; Wan et al., 2019; Wang et al., 2019a, 2019b; Yu et al., 2021). NLRs can function as singletons, in genetically linked pairs, or in immune networks (Adachi et al., 2019; Wu et al., 2018).

Paired NLRs, located adjacent to one another on the chromosome, are common in plant genomes and characterized as “sensor” or “executor” based on their role in immune activation. Sensor NLRs recognize effectors and then cooperate with executor NLRs to activate immune signaling (Adachi et al., 2019; Wu et al., 2018). A particularly interesting subset of paired NLRs are those sensors containing a non-canonical integrated domain (ID), exemplified by the TNL RRS1 (RESISTANT TO RALSTONIA SOLANACEARUM 1) /RPS4 (RESISTANT TO P. SYRINGAE 4) pair in Arabidopsis and the CNL RGA5/RGA4 (R-GENE ANALOG 5/4) pair in rice. Here, a WRKY transcription factor-domain ID, derived from one of a family major regulators of plant immune responses (Wani et al., 2021), is found in the RRS1 sensor, and a HMA (heavy metal-associated) ID is found in the RGA5 sensor (Cesari et al., 2014a; Césari et al., 2014b; Saucet et al., 2015; Sarris et al., 2015; Van de Weyer et al., 2019). In the resting state, the sensors, RRS1 or RGA5, maintain their respective pairs in an inactive state. Upon effector recognition, sensor and executor cooperate to transduce immune signaling (Cesari et al., 2014a; Césari et al., 2014b; Ma et al., 2018). However, it is unclear how paired NLRs without ID in their sensors are regulated.

The Arabidopsis *CSA1* (*CONSTITUTIVE SHADE-AVOIDANCE1*) and *CHS3* (*CHILLING SENSITIVE 3*) form a head-to-head gene pair on chromosome 5 with ∼3.9 kb separating their start codons (Xu et al., 2015; Figure S1A). *CSA1* encodes a typical TNL. *CHS3* encodes an atypical TNL containing an additional LIM (Lin-11, Isl-1 and Mec-3) and DA1-like (containing one LIM and a conserved C-terminal) ID at the C terminus (Faigón-Soverna et al., 2006; Yang et al., 2010; Figure S1A). *chs3-2D* is a gain-of-function mutant where a C1340Y substitution close to the LIM domain of CHS3 (Figure S1A), results in an auto-immune phenotype (Bi et al., 2011). CHS3 is the proposed sensor NLR and CSA1 is the proposed executor NLR in this pair (Van de Weyer et al., 2019). Arabidopsis allelic CHS3/CSA1 pairs are classified into three clades. Only clade 1 sensor CHS3 carries an ID domain (Van de Weyer et al., 2019; Figure 1A and S1B). Clade 1 CHS3 alleles share approximately 57% amino acid identity compared to clade 2 or clade 3, not including the ID; clade 2 and clade 3 CHS3 alleles share approximately 71% amino acid identity. Clade 1 CSA1 alleles share 62% amino acid identity compared to clade 2 or clade 3; clade 2 and clade 3 CSA1 alleles share approximately 86% amino acid identity. The co-formation of CHS3/CSA1 alleles in Arabidopsis into three matching clades suggests co-evolution (Van de Weyer et al., 2019), but there is no experimental data to support this hypothesis. And it is not known whether CHS3/CSA1 pairs from the three clades are differentially regulated, given their polymorphisms and the presence or absence of the ID in CHS3 alleles from clade 1. We suspected that since pre- and post-ID integration pair types co-exist, we might uncover differences in their activation.

**Figure 1.**
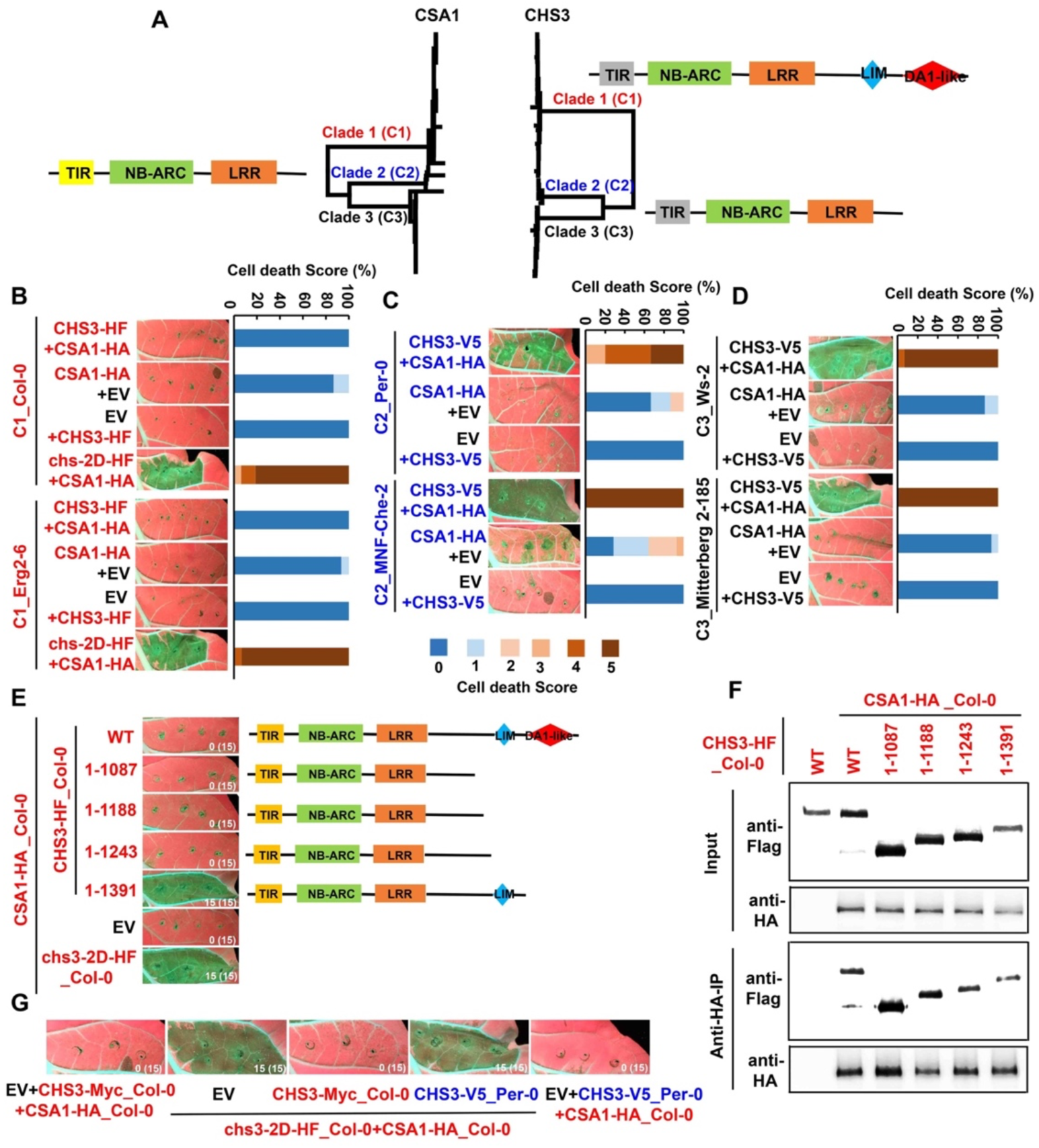
Wild-type CHS3/CSA1 pairs from clade 2 and clade 3, but not clade 1, trigger cell death, in *N. tabacum*. (A) Schematic phylogenetic tree of CSA1 (left) and CHS3 (right). C1, C2 and C3 represent clade 1, clade 2 and clade 3 respectively. (B-D) In planta (*N. tabacum*) phenotypes (left) and corresponding percentage representations of cell death score (right). Images were photographed at 4-5 dpi (days post infiltration). Clade 1 accessions and proteins are in red, clade 2 are in blue and clade 3 are in black, respectively. This color code is maintained throughout. HF: 6⊗His-36⊗Flag. EV: empty vector. Stacked bars are color-coded showing the proportions (in percentage) of each cell death score (0–5). 15 leaves were scored for each stacked bar. (E) Schematic of full-length and truncated clade 1 CHS3_Col-0 and in planta phenotypes. Co-infiltration of CSA1-HA_Col-0 with chs3-2D-HF_Col-0 is a positive control. EV: empty vector. Images were photographed at 4-5 dpi. The numbers indicate the numbers of leaves showing cell death out of the total number of leaves infiltrated. (F) Full-length and all truncated CHS3s associate with CSA1. Total proteins were extracted from *Nb* (*N. benthamiana*) at 2 dpi. Input: immunoblots showing the level of CSA1 and CHS3 in the total protein extract. Anti-HA-IP: immunoprecipitation products precipitated by the anti-HA antibody. Total proteins (input) or immunoprecipitation products are probed in immunoblots with antibodies to Flag or HA. (G) Cell death assays showing that clade 1 wild-type CHS3_Col-0 co-expression inhibits chs3-2D/CSA1-mediated cell death. Images were photographed at 4-5 dpi.

Here, we demonstrate that expression of CSA1 executor proteins alone cannot trigger visible cell death. By contrast, co-expression of intra- or inter-clade combinations of CSA1 with CHS3 alleles from clade 2 and clade 3 do induce cell death. This is not the case for clade 1, where the DA1-like ID contributes to maintenance of the clade 1 CHS3/CSA1 complex in an inactive state; its deletion leads to activation. Our data demonstrate functional co-evolution and specialization in allelic variants of the CHS3/CSA1 pair; clade 1 CHS3/CSA1 pairs are distinct from clades 2 and 3 because they cannot associate and function with each other in inter-clade combinations, even after ID deletion. Additionally, an intact p-loop in both CHS3 and CSA1 is required for complex formation and function across all clades, demonstrating that both sensor and executor contribute actively to function. We identified by natural variation two CSA1 residues that determine intra- and inter-clade allelic specialization. We additionally discovered that the LRR-kinase MAMP co-receptor BAK1 (BRI1-ASSOCIATED RECEPTOR KINASE), which plays a central role in multiple PTI signaling pathways, is guarded by CHS3/CSA1 pairs from all clades. The auto-immune phenotype of Arabidopsis mutants made by loss of function of *BAK1* and its closest paralog *BKK1* (*BAK1-LIKE 1*) in the Col-0 ecotype required both clade 1 CSA1 and CHS3. Consistent with this, BAK1 expression inhibits the cell death induced by clade 2 and 3 CHS3/CSA1 pairs, and previous data shows that the BAK1 signaling is targeted by an effector (Li et al., 2016; Wu et al., 2020). Together, our results suggest that the “guard and guardee” system is a conserved mechanism to regulate paired NLRs and further demonstrate that this mechanism can tolerate acquisition of an ID and a second regulatory mode.

## Results

### Intra-clade wild type CHS3/CSA1 pairs from clades 2 and 3, but not clade 1, trigger cell death

CHS3 and CSA1 alleles can be divided into three phylogenetic clades (Van de Weyer et al., 2019; Figure 1A and S1B). However, the functional relevance of this division is unclear. Ectopic expression of clade 1 CSA1 from accession Columbia-0 (Col-0) alone is not sufficient to induce evident cell death, but co-expression of clade1 Col-0 CSA1 with the chs3-2D gain-of-function mutant did result in cell death in *Nicotiana benthamiana* (*N. benthamiana*) (Castel et al., 2019). To investigate if expression of CSA1 alleles from other accessions and different clades is sufficient to trigger cell death, we constructed HA-tagged CSA1s (Figure S1B) and transiently expressed them in *Nicotiana tabacum* (*N. tabacum*) leaves. We found that none of these CSA1s triggered evident cell death in *N. tabacum* (Figure S1C and S1D). To examine if the sensor CHS3 is also required to activate the executor CSA1 and induce cell death, we transiently co-expressed several intra-clade CHS3/CSA1 pairs in *N. tabacum*. We re-capitulated previous findings (Castel et al., 2019) that co-expression of clade 1 CSA1 with chs3-2D but not with wild-type CHS3, elicited cell death (Figure 1B and S2A; cell death scale is defined in Figure S2B). By contrast, co-expression of intra-clade wild-type CSA1 and CHS3 from clade 2 or clade 3 triggered cell death in *N. tabacum* (Figure 1C, 1D and S2C).

Given the presence or absence of ID polymorphism in the CHS3 alleles (Figure 1A), we used truncated CHS3 derivatives from clade 1 Col-0 to test whether the integrated domain (ID) acts an inhibitor to maintain the clade 1 pair in its inactive state. We co-expressed CSA1 with either full-length or a series of truncated CHS3. We found that deleting the DA1-like ID from CHS3 resulted in cell death (Figure 1E), indicating the DA1-like ID contributes to maintaining the clade 1 CHS3/CSA1 pair in its inactive state. Co-immunoprecipitation (Co-IP) assays revealed that wild-type CHS3 and all truncated derivatives associated with CSA1 (Figure 1F), even in the absence of cell death. Moreover, we found that co-expression of wild-type CHS3 from clade 1 Col-0, but not that of clade 2 Per-0, abolished Col-0 chs3-2D/CSA1-mediated cell death (Figure 1G and S2D). Consistent with these data, *chs3-2D*-dependent autoimmunity is recessive (Figure S2E). The cell death induced by all intra-clade CHS3/CSA1 pairs were NRG1- and EDS1-dependent (Figure S2F-S2H). Weaker phenotypes were observed in *N. benthamiana* (Figure S2F), so we focused on *N. tabacum* throughout (Figure S2F). Taken together, we propose that paired sensors with ID regulate their corresponding executor NLRs (Cesari et al., 2014a; Césari et al., 2014b; Saucet et al., 2015; Sarris et al., 2015), whereas paired sensors without IDs do not inhibit their executors (Figure S2I). Consistent with this, expression of the TNL pair CHS1/SOC3 which does not carry an ID did trigger cell death (Figure S2J).

### Intact p-loops from both CHS3 and CSA1 are required for the complex formation and cell death induction

To further investigate the mechanistic requirements for CHS3/CSA1 pair cell death activation, we mutated both the CHS3 and CSA1 p-loop motifs. The p-loop is conserved in NLR proteins and typically is required for ATP binding and NLR oligomerization (Martin et al., 2020; Meyers et al., 1999; Jacob et al., 2021; Saraste et al., 1990; Walker et al., 1982; Wang et al., 2019a). Here, we selected one CHS3/CSA1 pair from each clade (clade 1 Col-0, clade 2 Per-0 and clade 3 Ws-2) to test if the p-loop is required for cell death induction. For clade1 Col-0, we used chs3-2D (Figure 1B). Putative catalytic dead p-loop alleles from either CHS3 or CSA1 abrogated the cell death responses triggered by these three CHS3/CSA1 pairs (Figure 2A). Additionally, co-immunoprecipitation (Co-IP) assays revealed that p-loop mutation in CHS3 or CSA1 reduced their association (Figure 2B-2D), suggesting that an intact p-loop in both CHS3 and CSA1 is necessary for complex formation and cell death induction in all clades. These results indicated that the function of the p-loop in CSA1 and CHS3 is conserved across all clades and that both functional sensor and executor are essential to execute cell death. These results suggest that the sensor NLR in some pairs contributes actively to function and is not merely a negative regulator of otherwise constitutive executor function.

**Figure 2.**
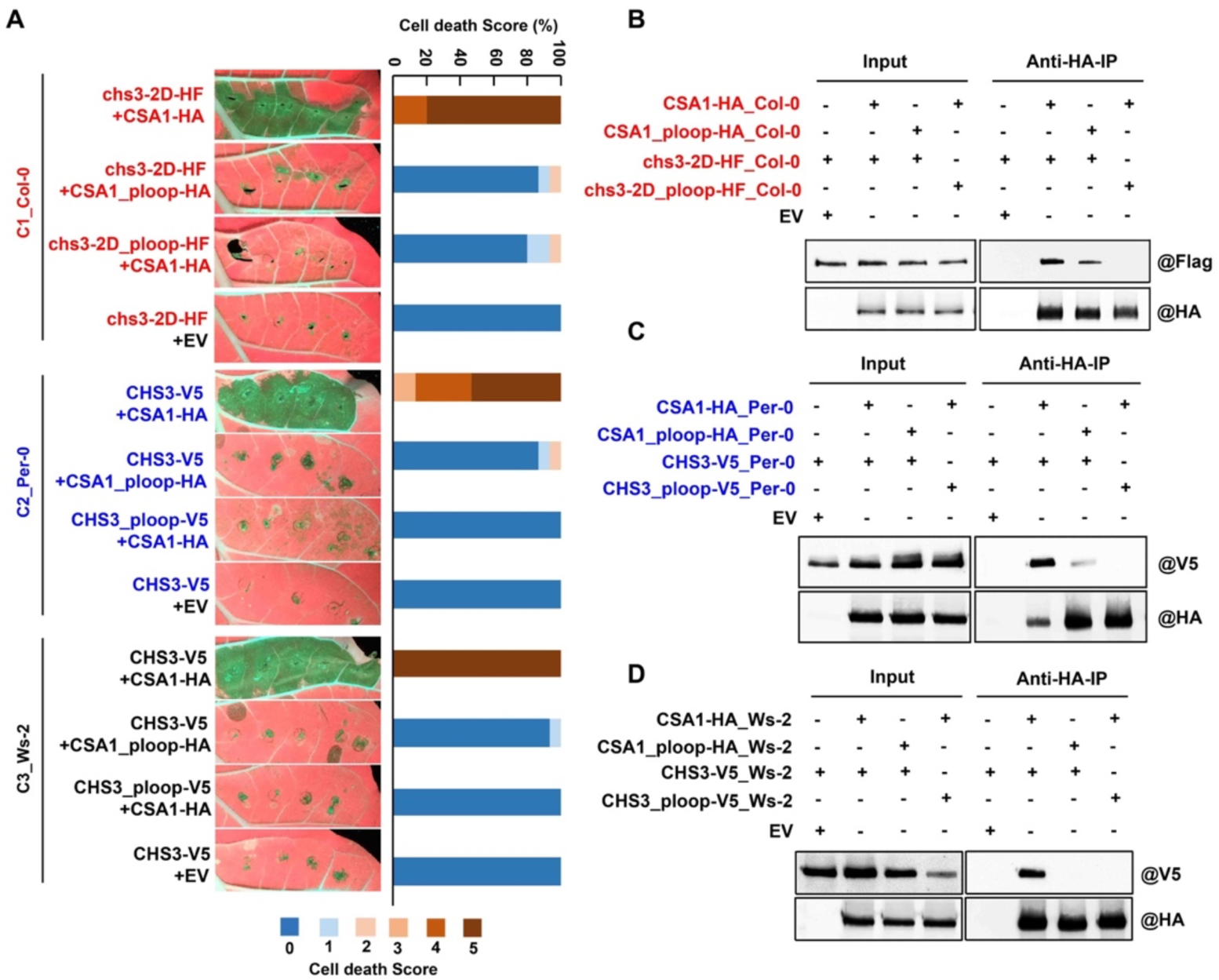
Intact p-loops of CHS3 and CSA1 are required for cell death and association. (A) In planta (*N. tabacum*) phenotypes (left) and corresponding percentage representations of cell death score (right). Clade 1 accessions and proteins are in red, clade 2 are in blue and clade 3 are in black. Images were photographed at 4-5 dpi. EV: empty vector. Stacked bars are color-coded showing the proportions (in percentage) of each cell death score (0–5). 15 leaves were scored for each stacked bar. (B-D) P-loop dead alleles of CHS31 or CSA1 exhibit decreased association. Total proteins were extracted from *N. benthamiana* at 2 dpi. Input: immunoblots showing the level of CSA1 and CHS3 in the total protein extract. Anti-HA-IP: immunoprecipitation products precipitated by the anti-HA antibody.

### Alleles from different clades of the CHS3/CSA1 TNL pair exhibit functional co-evolution and specialization

To analyze functional specificity in allelic CHS3/CSA1 pairs, we co-expressed different intra-or inter-clade CHS3/CSA1 combinations and assessed their ability to trigger cell death in *N. tabacum*. Co-expression of the clade 1 chs3-2D_Col-0 with clade 1 CSA1 from Col-0, Erg2-6 or Scm-0 led to strong cell death (Figure 3A). However, we observed no cell death after co-expression of clade 1 chs3-2D_Col-0 with CSA1 from clade 2 or clade 3 (Figure 3A, S3A and S3B). As expected, substitution of chs3-2D with wild-type CHS3 did not result in cell death (Figure S3C). Conversely, co-expression of inter-clade combinations of clade 1 CSA1 with clade 2 CHS3_Per-0 or clade 3 CHS3_Bar-1 also did not result in cell death (Figure 3A). Thus, inter-clade pairs consisting of clade 1 CSA1 or CHS3 together with CHS3 or CSA1 from clade 2 or clade 3 are not functional in this cell death assay. These data suggest intra-clade co-evolution of CHS3 (with its ID domain) and CSA1 from clade 1.

**Figure 3.**
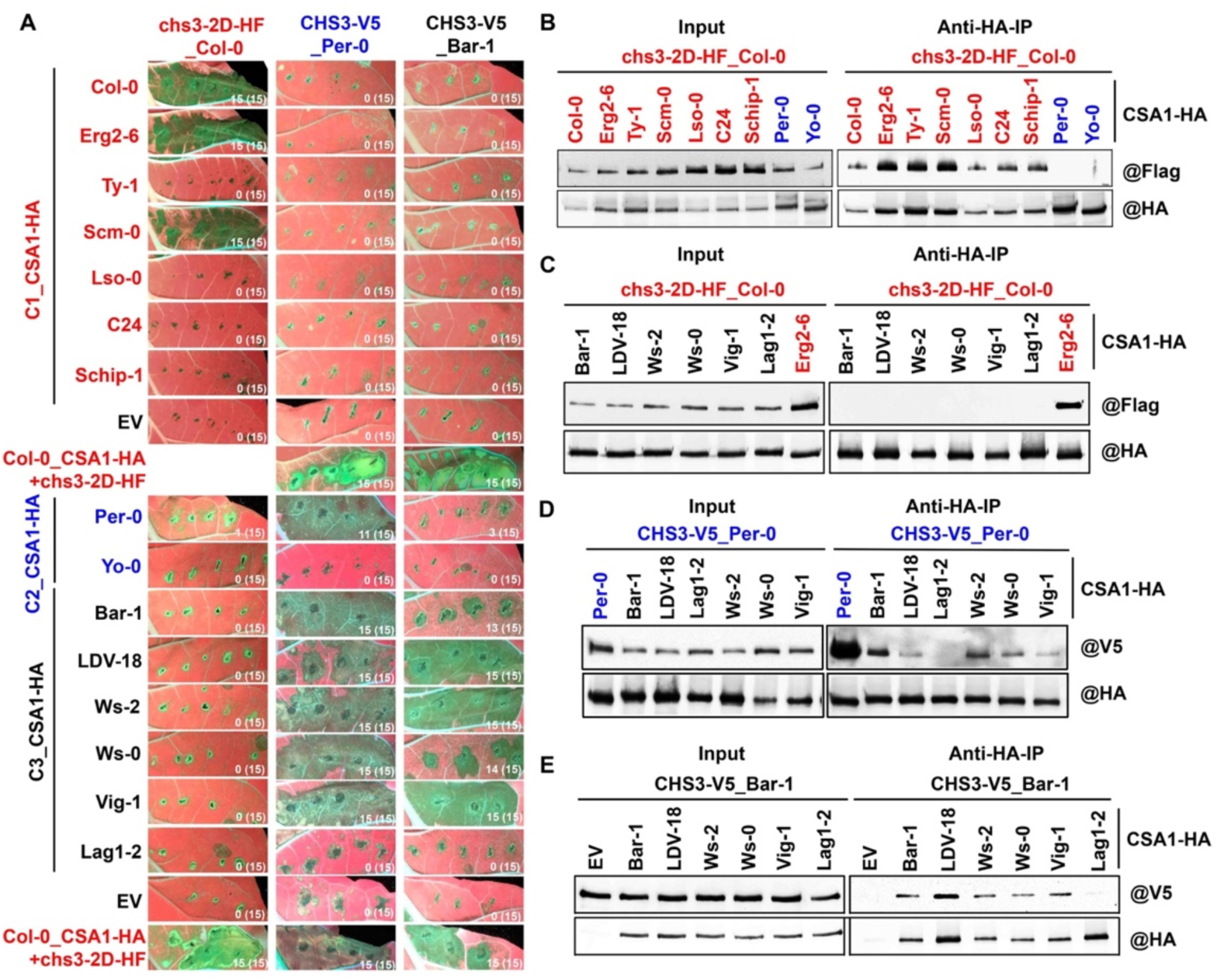
Inter-clade combinations of clade 1 CSA1 or chs3-2D with clade 2 or clade 3 CHS3 or CSA1 do not induce cell death or association, while functional inter-clade combinations between clade 2 and clade 3 do. (A) In planta phenotypes. Co-infiltration of Col-0 CSA1-HA with chs3-2D-HF is a positive control. Images were photographed at 4-5 dpi. EV: empty vector. Clade 1 accessions and proteins are in red, clade 2 are in blue and clade 3 are in black. The numbers indicate the numbers of leaves displaying cell death out of the total number of leaves infiltrated. (B and C) Clade 1 HF-tagged chs3-2D_Col-0 associates with all clade 1 HA-tagged CSA1s (B), but not with clade 2 and clade 3 CSA1s (B and C). Total proteins were extracted from *N. benthamiana* at 2 dpi. Clade 1 accessions and proteins are in red, clade 2 are in blue and clade 3 are in black. (D and E) Association of intra- and inter-clade 2 and 3 CHS3/CSA1 combinations. Clade 2 V5-tagged CHS3_Per-0 associates with HA-tagged clade 2 CSA1_Per-0 and all tested clade 3 CSA1s of except CSA1_Lag1-2(D). Clade 3 V5-tagged CHS3_Bar-1 associates with all clade 3 HA-tagged CSA1s except CSA1_Lag1-2 (E). clade 2 accessions and proteins are in blue and clade 3 are in black.

By contrast, inter-clade combinations from clade 2 and clade 3 were typically functional. When we co-expressed clade 2 CHS3_Per-0 with clade 3 CSA1, we observed cell death in *N. tabacum*, except in the case of clade 3 CSA1_Lag1-2. Similarly, intra-clade co-expression of clade 3 CHS3_Bar-1 with most clade 3 CSA1s resulted in cell death. Again, the exception was the CSA1_Lag1-2. Meanwhile, the intra-clade combinations of clade 3 CHS3_Bar-1 with CSA1_Bar-1 or CSA1_Ws-0 induced weak cell death (Figure 3A). All the combinations we tested were expressed (Figure S3A and S3B). As a control, the CSA1 allele from accession Yo-0 is highly likely to lack the NADase activity encoded in the TIR domain due to mutation of the catalytic glutamic acid, so it cannot trigger cell death following co-expression with any CHS3s. This last result suggests that TIR NADase activity is required for CHS3/CSA1 pair cell death.

To determine if the differential cell death phenotypes resulted from the association of the tested allelic CHS3/CSA1 combinations, we performed co-immunoprecipitation (Co-IP) assays. All clade 1 CSA1s that we tested associated with chs3-2D (cell death observed) and wild-type CHS3 (no cell death) (Figure 3B and S3D). However, inter-clade combinations of chs3-2D or CSA1 from clade 1 did not associate with CSA1 or CHS3, respectively, from clade 2 or clade 3, consistent with the lack of cell death (Figure 3B, 3C, S3E and S3F). Additionally, Co-IP assays demonstrated that functional intra-or inter-clade combinations from clade 2 and clade 3 did associate, but combinations containing the non-functional CSA1_Lag1-2 did not (Figure 3D and 3E).

Given that the DA1-like ID truncation of clade 1 CHS3_Col-0 activates CSA1_Col-0 and triggers cell death (Figure 1E), we examined whether the truncated CHS3_Col-0 could also support inter-clade activation of CSA1s. We transiently co-expressed the truncated clade 1 CHS3_Col-0 with CSA1 from clade 1 or clade 3. Note that the clade 3 CSA1s we used supported strong inter and intra-clade cell death when co-infiltrated with CHS3 of clade 2 or clade 3 (Figure 3A). We observed that the intra-clade combination of the clade 1 DA1-like truncation of CHS3 with clade 1 CSA1_Erg2-6 led to cell death, but that no cell death was observed in the inter-clade combinations with clade 3 CSA1 (Figure S4A and S4B). The cell death phenotype observed for all intra- and inter-clade CHS3/CSA1 pairs was also dependent on NRG1 and EDS1, and on intact p-loops of both CHS3 and CSA1 (Figure S4C-S4E).

Altogether, we demonstrate that CHS3/CSA1 pairs from different clades have co-evolved and diverged in their requirements for association and regulation. Clade 1 CHS3 or CSA1 cannot associate and function with clade 2 or clade 3, even if the DA1-like ID is deleted from the clade 1 CHS3. Additionally, we found that most of the differential cell death phenotypes induced by intra- and inter-clade 2 and clade 3 combination of CHS3/CSA1 pairs were correlated with alterations in association. This is in contrast to the intra-clade CHS3/CSA1 combinations from clade 1, where association is independent of cell death induction.

### A conserved glycine residue determines allelic specialization and is essential for CHS3/CSA1 pair function in all clades

To identify structural specificities underpinning the differential cell death phenotype triggered by intra- and inter-clade CHS3/CSA1 combinations, we aligned the full-length protein sequences from both functional and non-functional clade 1 CSA1s or clade 3 CSA1s that were tested in Figure 3. We did not identify any residues that strictly correlated with cell death induction from the seven clade 1 sequences. By contrast, we identified a single amino acid in clade 3 CSA1s. In the cell death inactive CSA1_Lag1-2 allele, the glycine (G) 681, which is conserved in the cell death-inducing clade 3 CSA1s, is replaced by a glutamic acid (E) residue (Figure S5A). There are other different residues between the clade 3 CSA1s but only G681E was strictly correlated with cell death induction. To determine if CSA1 G681 is required for cell death induction, we generated G681E and E681G CSA1s from clade 3 and co-expressed these with clade 2 CHS3_Per-0 or clade 3 CHS3_Bar-1. While wild-type CSA1_Lag1-2 fails to induce cell death, E-to-G mutation in clade 3 CSA1_Lag1-2 enhanced the cell death phenotype in combination with clade 2 CHS3_Per-0 or clade 3 CHS3_Bar-1 (Figure 4A and S5B-S5D). Conversely, the cell death phenotype of the clade 3 CSA1 G681E mutants was decreased when co-expressed with clade 2 CSA1_Per-0 or clade 3 CHS3_Bar-1 (Figure 4A and S5B-S5D). The ability of clade 3 CSA1_Lag1-2 E681G to trigger cell death in combination with clade 2 CHS3_Per-0 or clade 3 CHS3_Bar-1 was surprising given that wild-type proteins do not interact in co-IP, even though other functional inter-or intra-clade pairs do associate (Figure 3D and 3E). We tested if G681 affects the ability of clade 3 CSA1s to form hetero-complexes with clade 2 CHS3_Per-0. Consistent with the cell death phenotypes, G-to-E mutation in clade 3 CSA1s diminished their association, while E-to-G mutation in clade 3 CSA1_Lag1-2 enhanced association with clade 2 CHS3_Per-0 (Figure 4B). These results demonstrate that G681, or equivalent residues in clade 3 CSA1s, is essential for cell death induction and pair association.

**Figure 4.**
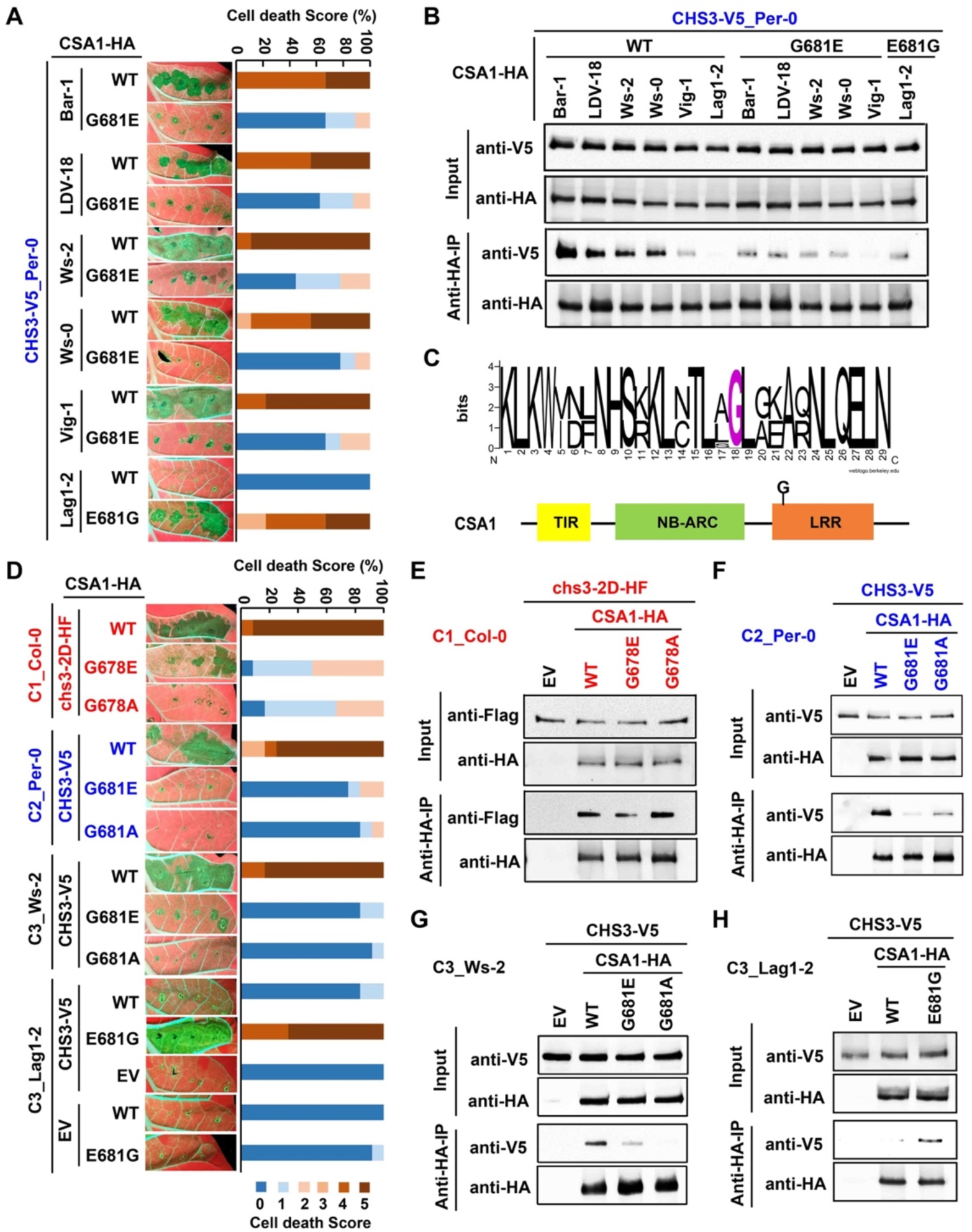
A single glycine residue in CSA1 is conserved in all clades and is essential for CHS3/CSA1 function. (A) CSA1 G681 is required for cell death induction by inter-clade combinations of clade 2 CHS3_Per-0 with clade 3 CSA1s. Images were photographed at 4-5 dpi. Amino acid abbreviations: G, glycine; E, glutamic acid. Clade 2 accessions and proteins are in blue and clade 3 are in black. Stacked bars are color-coded showing the proportions (in percentage) of each cell death score (0–5). 9 leaves were scored for each stacked bar. (B) Co-IP assays showing clade 3 CSA1 G681 is essential for association with clade 2 CHS3_Per-0. Clade 2 accessions and proteins are in blue and clade 3 are in black. (C)A sequence logo showing conserved G residue in LRR domain of CSA1s from 60 Arabidopsis accessions and schematic representation of G residue position. (D) In planta phenotypes (left) and corresponding percentage representations of cell death scores (right). EV: empty vector. Clade 1 accessions and proteins are in red and clade 2 are in blue and clade 3 are in black. Stacked bars are color-cocoded and show the proportions (in percentage) of each cell death score (0–5). 12 leaves were scored for each stacked bar. (E-H) Co-IP assays for association of matched CHS3/CSA1 pairs with or without G to E/A or E to G mutation in CSA1s. EV: empty vector. Clade 1 accessions and proteins are in red, clade 2 are in blue and clade 3 are in black.

We aligned all full-length CSA1 amino-acid sequences and found that the G residue is conserved in all clades (Figure 4C). Therefore, we introduced the G-to-E mutation at the equivalent position 678 in clade 1 CSAs; this reduced cell death activity of chs3-2D/CSA1 combinations (Figure S5B-S5D). To test whether CSA1 G678/681 residue is indispensable for cell death mediated by intra-clade matched pairs, we generated potential loss-of-function mutation with either G to E or G to A (alanine) in clade 1 CSA1_Col-0, clade 2 CSA1_Per-0 and clade 3 CSA1_Ws-2 and potential gain-of-function mutation with E to G in clade 3 CSA1_Lag1-2. We co-expressed either each wild-type CSA1 or each mutant CSA1 with their corresponding CHS3. We found that G-to-E or G-to-A mutation reduced all matched pairs-mediated cell death (Figure 4D). By contrast, E-to-G mutation increased the cell death phenotype induced by clade 3 Lag1-2 CHS3/CSA1 pair (Figure 4D).

We also confirmed that the G residue also affects association of intra-clade pairs. Interestingly, the G residue had minor effects on association of the clade 1 Col-0 CHS3/CSA1 pair (Figure 4E) but was required for association of intra-clade matched pairs from clade 2 and clade 3 (Figure 4F-4H). Analysis of 74 *Arabidopsis* TNLs from the Col-0 reference genome revealed that this residue is conserved in all RPS4 - like TNLs (analogous to CSA1) (Figure S5E). To further validate its function, we generated the relevant G-to-E mutation in RPS4. Compared to wild-type RPS4, this mutant did not trigger cell death in *N. benthamiana* leaves when it was co-expressed with RRS1 and AvrRps4 (Figure S5F).

### Another single amino acid specific to clade 2 and clade 3 CSA1s is required for their cell death induction

We next investigated why co-expression of the clade 3 pair of CHS3_Bar-1 with CSA1_Ws-0 directed weaker cell death than other clade 3 pairs (Figure 3A). We aligned full-length protein sequences from the clade 3 CSA1s tested in Figure 3A. We found that CSA1_Bar-1 and CSA1_Ws-0 differed from other clade 3 CSA1s at position 543, where aspartic acid (D) was replaced by an asparagine (N) (Figure S5G). Since the matched CHS3/CSA1 pair from Bar-1 induced weaker cell death as well, we tested whether the CSA1 N543 residue underpinned the weaker cell death phenotype. We introduced the aspartic acid (D)-to-asparagine (N) and N-to-D mutations in clade 3 CSA1s. We observed that the D-to-N mutation decreased cell death; conversely the N-to-D mutation enhanced cell death in intra-clade pairs from clade 3 (Figure 5A and S5H).

**Figure 5.**
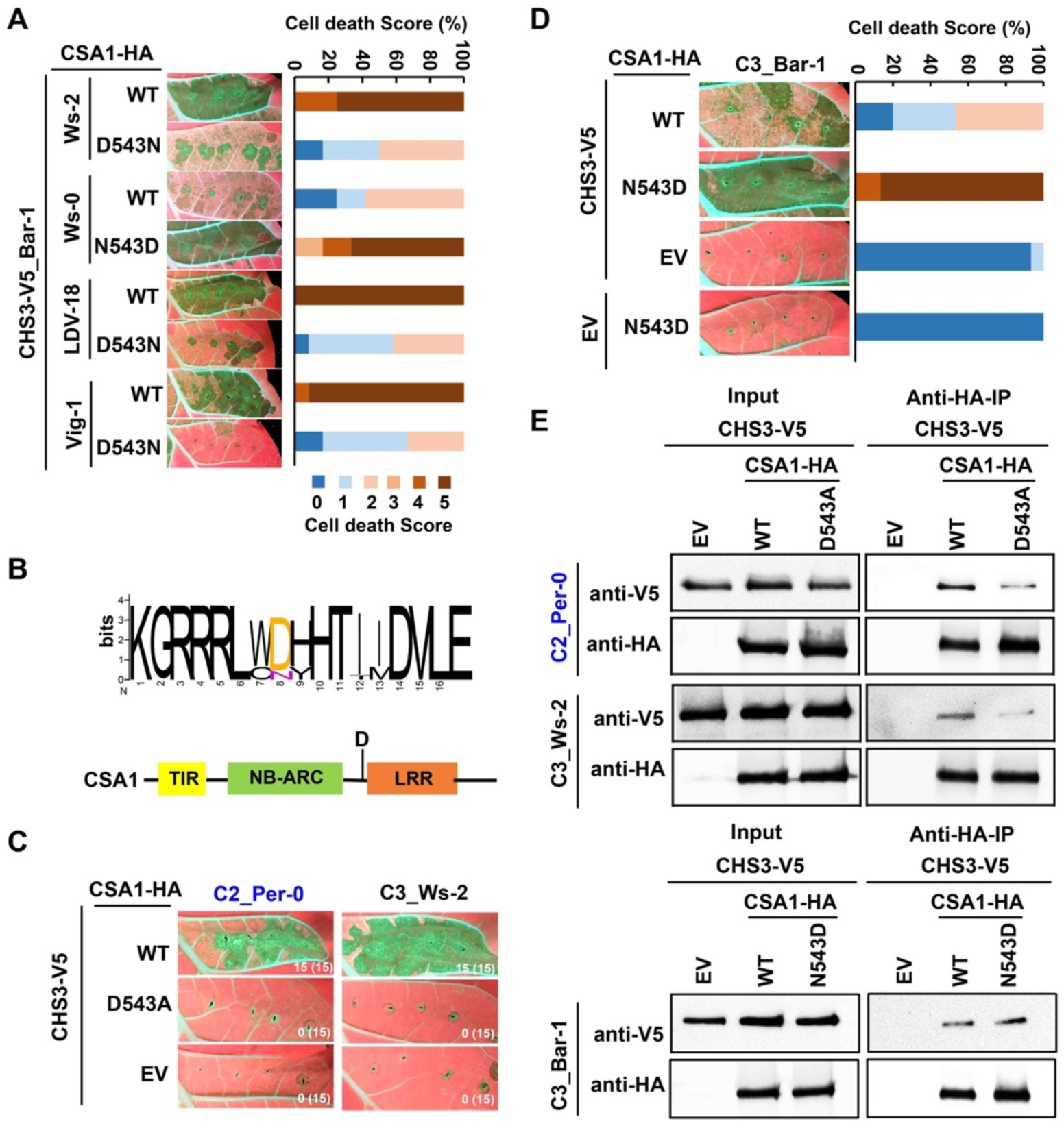
D543 is conserved only in clade 2 and clade 3 CSA1s and is required for CHS3/CSA1 function and association. (A) In planta phenotypes (left) and corresponding percentage representations of cell death scores (right). Images were photographed at 4-5 dpi. Amino acid abbreviations: D, Aspartic acid; N, Asparagine. Stacked bars are color-coded showing the proportions (in percentage) of each cell death score (0–5). 12 leaves were scored for each stacked bar. (B) Sequence pileup of the CSA1s from clade 2 and clade 3 and schematic representation of the D residue position. (C) Effect of D543A mutation on cell death induction by clade 2 and clade 3 matched CHS3/CSA1 pairs. Images were photographed at 4-5 dpi. The numbers represent the numbers of leaves with cell death out of the total number of leaves infiltrated. (D) Effect of N543D mutation on cell death induction (left) and corresponding percentage representations of cell death scores (right). Images were photographed at 4-5 dpi. 15 leaves were scored for each stacked bar. (E) Co-IP assays assess association of matched CHS3/CSA1 pairs with or without D543A or N543D mutation in the respective CSA1s. Accessions from clade 2 are in blue and clade 3 are in black. (D) In planta phenotypes (left) and corresponding percentage representations of cell death scores (right). EV: empty
vector. Clade 1 accessions and proteins are in red and clade 2 are in blue and clade 3 are in black. Stacked bars are
color-coded and show the proportions (in percentage) of each cell death score (0–5). 12 leaves were scored for each stacked bar. (E-H) Co-IP assays for association of matched CHS3/CSA1 pairs with or without G to E/A or E to G mutation in CSA1s. EV: empty vector. Clade 1 accessions and proteins are in red, clade 2 are in blue and clade 3 are in black.

Alignment of CSA1 protein sequences from all clades revealed that this D residue is conserved only in clade 2 and clade 3 (Figure 5B). We assessed the functional relevance of this D residue in matched pairs from clade 2 Per-0 and clade 3 Ws-2, and we found that D-to-A mutation decreased their cell death phenotypes (Figure 5C). We generated the N-to-D mutation in clade 3 CSA1_Bar-1 and found that this increased cell death in combination with clade 3 CHS3_Bar-1 (Figure 5D). There are only two residues difference between the full length CSA1_Ws-2 and CSA1_Ws-0. Co-IP assays showed there is stronger intra-clade association of clade 3 CHS3_Bar-1 with CSA1_Ws-2 than with CSA1_Ws-0 (Figure 3E and S5I). To examine whether D543 was responsible for this difference in protein association, we performed Co-IP assays and found that the D-to-A mutation reduced the association of clade 2 and clade 3 matching pairs, conversely, the N-to-D mutation enhanced the association of the clade 3 Bar-1 CHS3/CSA1 pair (Figure 5E). This aspartic acid is also conserved in the paired TNL SOC3, but it is not necessary for CHS1/SOC3-mediated cell death (Figure S5J and S5K). Additionally, we note that the conserved G residue described above is indispensable because clade 3 CSA1_Lag1-2 cannot support cell death induction despite the presence of the D543 residue (Figure S5A and S5G).

### Two distinct regulatory mechanisms maintain the CHS3/CSA1 pair in the inactive state

The clade 1 sensor CHS3 contains an ID that acts as a negative regulator to maintain clade 1 CHS3/CSA1 pairs in a resting state. There is no ID in CHS3s from clade 2 or clade 3 and transient co-expression of intra-or inter-clade CHS3/CSA1 pairs does result in cell death. Nevertheless, wild-type Arabidopsis accessions expressing these genes do not exhibit autoimmunity, suggesting that the resting state complexes containing clade 2 and clade 3 CHS3/CSA1 pairs are negatively regulated in the absence of pathogen. We speculated that other components are required to maintain CHS3/CSA1 pairs in a resting state. The receptor protein kinase BAK1 belongs to a five-member leucine-rich-repeat receptor kinase subfamily and as a co-receptor it plays a central role in multiple PTI signaling pathways. The mutants made in *BAK1* and its closest paralog *BKK1*, in the Col-0 background, exhibit helper NLR (ADR1 family)-dependent auto-immune phenotypes (He et al., 2007, 2008; Gao et al., 2017; Wu et al., 2020). Furthermore, the pathogen HopB1 virulence effector cleaves BAK1 and its paralogs (Li et al., 2016), leading to ADR1 family-dependent cell death (Li et al., 2016; Wu et al., 2020). These data suggested that BAK1 and its paralogs were ‘guarded’ by unknown NLRs that required helper NLR function. We hypothesized that the CHS3/CSA1 pair, whose activity depends on helper NLRs, contributed to the *bak1 bkk1* auto-immune phenotype.

We thus constructed the triple mutants of *csa1 bak1-3 bkk1-1* and *chs3 bak1-3 bkk1-1* in the Col-0 background using CRISPR/CAS9 (Figure S6A-S6C). Compared to *bak1-3 bkk1-1*, the auto-immune phenotype was largely suppressed in *csa1 bak1-3 bkk1-1* and partially suppressed in *chs3 bak1-3 bkk1-1* (Figure 6A and S6D). Consistent with these phenotypes, the increased *PR1* gene expression typical of auto-immune mutants was suppressed in *csa1 bak1-3 bkk1-1* and *chs3 bak1-3 bkk1-1* (Figure 6B and S6E). These data are consistent with the hypothesis that CHS3/CSA1 pair guards BAK1 and BKK1. This was further validated by the observation that *bak1-4* mutation enhanced the auto-immune phenotype of heterozygous *chs3-2D* (Figure 6C), as evidenced by increased *PR1* expression (Figure 6D). Co-IP assays demonstrated that both Col-0 BAK1 and BKK1 can associate with CSA1 and CHS3, respectively (Figure 6E and 6F). These results indicate that there are two distinct regulatory modes for clade 1 CHS3/CSA1 pairs. The first is the DA1-like ID-dependent negative regulation of clade 1 sensor CHS3 on the CHS3/CSA1 complex, detailed above, and the second mechanism is mediated by the “guardees” BAK1 and BKK1 which maintain clade 1 CHS3/CSA1 pairs in an inactive resting state.

**Figure 6.**
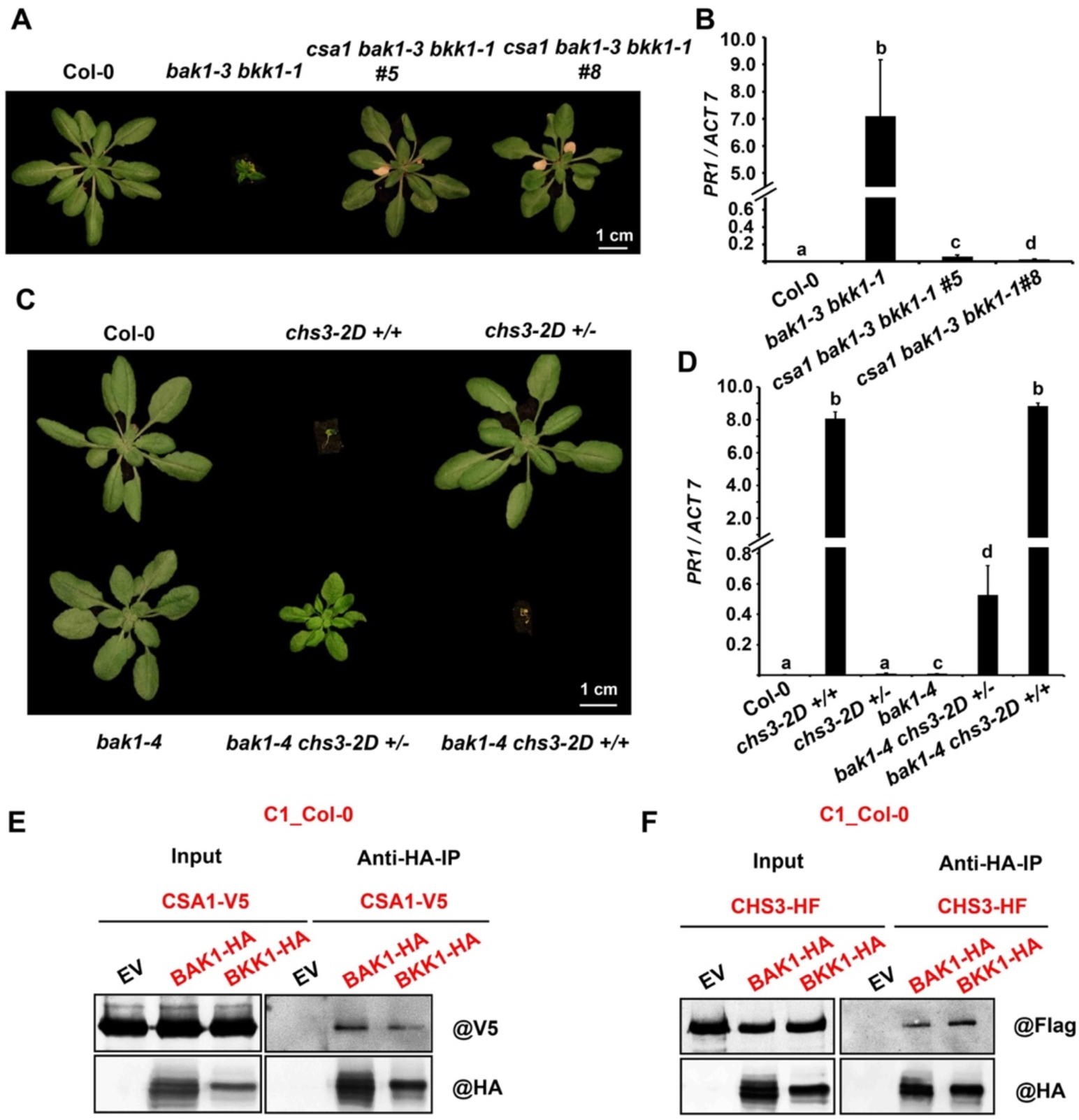
Clade 1 CSA1_Col-0 is required for the *bak1-3 bkk1-1* auto-immune phenotype. (A) Morphological phenotypes of indicated genotypes. *csa1* mutation largely suppresses *bak1-3 bkk1-1*-auto-immune phenotype. Plants were grown in 21°C/18°C under LD (16 h light/8 h dark) condition. Images were photographed at 3-5 weeks. (B) *PR1* gene expression in the indicated plants from (A) as determined by RT-PCR and normalized to *ACTIN7* (*ACT7*). Error bars represent two biological repeats (three technical replicates were performed for each biological repeat). The different letters “a-d” indicate statistically significant differences determined by one-way ANOVA for multiple pairwise comparisons and Tukey’s least significant difference test. P<0.05. (C) Morphological phenotypes of indicated genotypes are shown. *bak1* mutation enhances the auto-immune phenotypes of heterozygous *chs3-2D*. Plants were grown in 21°C/18°C under LD (16 h light/8 h dark) condition. Images were photographed at 3-5 weeks. +/+: homozygous *chs3-2D*, +/-: heterozygous *chs3-2D*. (D) *PR1* gene expression in the indicated plants from (C) as determined by RT-PCR and normalized to *ACTIN7* (*ACT7*). Error bars represent two biological repeats (three technical replicates were performed for each biological repeat). The different letters “a-d” indicate statistically significant differences determined by one-way ANOVA for multiple pairwise comparisons and Tukey’s least significant difference test. P<0.05. (E and F) Clade 1 Col-0 CSA1 (E) and CHS3 (F) can associate with corresponding BAK1 and BKK1. Total proteins were extracted from *N. benthamiana* at 2 dpi.

BAK1 is highly conserved in different Arabidopsis accessions. We next explored if association of BAK1 by CHS3/CSA1 pairs is conserved in clade 2 and clade 3. We co-expressed BAK1 with CSA1 or CHS3 from clade 2 Per-0 or clade 3 Ws-2 and performed Co-IPs. We found that BAK1 associated with CSA1 from Per-0 or Ws-2, but we did not detect association between BAK1 and CHS3 from Per-0 (Figure 7A and S7A). Absence or over-expression of BAK1 leads to deregulated cell death in Col-0 (Dominguez-Ferreras et al., 2015; He et al., 2007; Kemmerling et al., 2007). However, the *bak1-5* missense mutation in Col-0 reduces PTI responses and *bak1-5 bkk1* does not trigger auto-immunity (Schwessinger et al., 2011). To investigate if BAK1 or bak1-5 could inhibit cell death mediated by clade 2 or clade 3 CHS3/CSA1 pairs, we co-infiltrated constructs to express either BAK1 or bak1-5 with CHS3/CSA1 pairs from clade 2 Per-0 or clade 3 Ws-2. BAK1 alone induced weak and inconsistent cell death in *N. tabacum* (Figure 7B). We discovered that both BAK1 and bak1-5 inhibited the cell death induction by CHS3/CSA1 pairs from Per-0 or Ws-2 and we noted that the suppression of CHS3/CSA1 cell death by bak1-5 was stronger than wild-type BAK1 (Figure 7B and S7B). These data demonstrate that BAK1 negatively regulates cell death mediated by clade 2 or clade 3 CHS3/CSA1 pairs.

**Figure 7.**
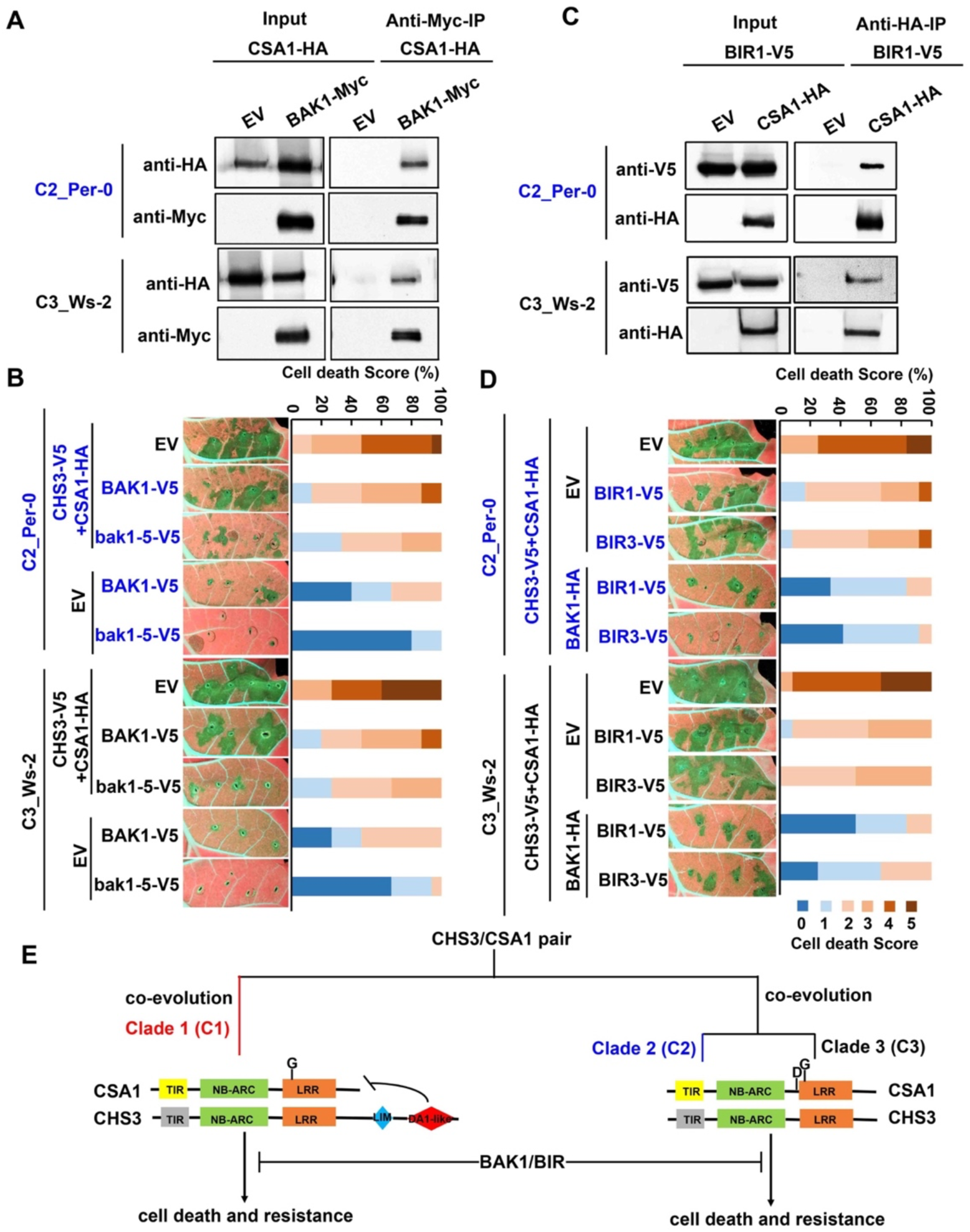
BAK1 associates with clade 2 CSA1_Per-0 and clade 3 CSA1_Ws-2 and suppresses cell death induced by clade 2 and clade 3 CHS3/CSA1 pairs. (A) Co-IP assays assess association of BAK1 with CSA1 from clade 2 or clade 3. Total proteins were extracted from *N. benthamiana* at 2 dpi. Accession from clade 2 is in blue and clade 3 is in black. (B) BAK1 inhibits cell death triggered by CHS3/CSA1 pairs from clade 2 Per-0 and clade 3 Ws-2. Clade 2 accessions and proteins show in blue and clade 3 show in black. Stacked bars are color-coded showing the proportions (in percentage) of each cell death score (0–5). 15 leaves were scored for each stacked bar. (C) BIR1 associates with clade 2 CSA1_Per-0 and clade 3 CSA1_Ws-2. Total proteins were extracted from *N. benthamiana* at 2 dpi. Clade 2 accession Per-0 shows in blue and clade 3 Ws-2 shows in black. (D) BIR1 or BIR3 alone weakly suppress, and with BAK1 strongly suppress, clade 2 and clade 3 CHS3/CSA1-mediated cell death. Images were photographed at 4-5 dpi. Clade 2 accessions and proteins are in blue and clade 3 are in black. Stacked bars are color-coded showing the proportions (in percentage) of each cell death score (0–5). 12 leaves were scored for each stacked bar. (E) Schematic representation of the evolutionary and functional model for the structurally diverse CHS3/CSA1 pairs.

CSA1 is required for auto-immune cell death of *bak1 bir3* in Arabidopsis Col-0 and clade 1 CSA1_Col-0 interacts directly with BIR3 and indirectly with BAK1 (Schulze et al., 2021). We therefore cloned BIR1 and BIR3 from clade 2 Per-0 and clade 3 Ws-2 and tested their association with CSA1 and CHS3. Co-IP assays showed that BIRs associated with CSA1 from clade 2 Per-0 and clade 3 Ws-2, respectively, but weak or no association was detected between BIRs and the corresponding CHS3s (Figure 7C and S7C-S7G). BIR1 or BIR3 co-expression only slightly decreased the cell death phenotype triggered by CHS3/CSA1 pairs compared to the combination of BIRs and the strong suppression mediated by BAK1 (Figure 7D and S7H). Overall, there are two distinct regulatory modes for CHS3/CSA1 pairs. Sensor-mediated negative regulation in the absence of pathogen is relevant only for clade 1 CHS3/CSA1 pairs with ID in the sensor; however, surveillance of BAK1/BIR homeostasis by CHS3/CSA1 pairs is conserved across all clades (Figure 7I)

## Discussion

### Regulation of paired NLRs by their “Sensor”

The ID in the clade 1 sensor CHS3 is required to keep CHS3/CSA1 pair in an inactive state. This is reminiscent of the TNL RRS1/RPS4 pair in Arabidopsis and the CNL RGA5/RGA4 pair in rice (Césari et al., 2014b; Ma et al., 2018). Executors RPS4 and RGA4 are typically regarded as autonomous cell death inducers, and these functions can be suppressed by their respective sensors RRS1 and RGA5 in the absence of pathogen (Césari et al., 2014b; Huh et al., 2017; Ma et al., 2018; Wirthmueller et al., 2007). However, our data suggest that clade 2 and clade 3 CHS3/CSA1 pairs do not function according to this model. CSA1s from all clades are unable to trigger evident cell death when expressed alone, but co-expression of clade 2 or clade 3 CHS3 and CSA1 together does (Figure 1C and 1D). Moreover, intact p-loops from both CHS3 and CSA1 are required for cell death induction and their association in all clades (Figure 2). The p-loop dependence of both clade 1 CHS3_Col-0 and CSA1_Col-0 in chs3-2D/CSA1-mediated cell death was independently confirmed (Parkes, 2020). These results imply that CSA1 must associate with CHS3 and that both must maintain p-loop function to form a functional executor cell death initiation complex.

Other functional studies of the rice Pik pairs (including the Pikp pair and the Pikm pair), the Pias pair and the *Brassica napus Bn*RPR2/*Bn*RPR1 pair revealed that the sensors are indispensable for paired NLRs to trigger strong cell death, even though these sensors contain IDs (De la Concepcion et al., 2021; Mermigka et al., 2021; Shimizu et al., 2021; Zdrzałek et al., 2020). Moreover, an intact p-loop in the sensor of the Pik pair is required to recognize effector and elicit cell death (De la Concepcion et al., 2021; Zdrzałek et al., 2020). Previous papers reported that the p-loop in the executors RPS4 and RGA4, but not the respective sensors RRS1 and RGA5, is required for effector-mediated cell death (Césari et al., 2014b, Williams et al., 2014). Recently, however, an intact p-loop and phosphorylation of the sensor RRS1 was shown to be required for auto-active RPS4 alleles to elicit strong cell death (Guo et al., 2021). Thus, both sensor and executor NLRs can be active participants in paired NLR signaling.

### Experimental data support intra-clade co-evolution of CHS3 and CSA1

CSA1 and CHS3 alleles are divided into three matching sets of phylogenetic clades based on nucleotide or full-length protein sequence (Van de Weyer et al., 2019; Figure 1A and S1B). High and co-incident Tajima’s D values of CSA1 and CHS3 suggested their co-evolution (Tajima, 1998; Van de Weyer et al., 2019). Our experimental data support the theory of diverging co-evolution of CSA1 and CHS3 clades. Wild-type matched CHS3/CSA1 pairs from clade 2 and clade 3, but not clade 1, trigger cell death response in *N. tabacum* or *N. benthamiana* leaves (Figure 7I). Additionally, inter-clade combinations from clade 2 and clade 3 also trigger cell death (Figure 3A). However, clade 1 CSA1 or CHS3 cannot cooperate with clade 2 or clade 3 proteins to cause cell death (Figure 3A). A truncated clade1 CHS3 (1-1391) lacking the DA1-like ID triggers cell death in intra-clade combinations with clade 1 CSA1s but cannot induce cell death in inter-clade combinations with clade 3 CSA1s (Figure 1E, S4A and S4B). Additionally, the clade 1 CHS3/CSA1 pair in Col-0 confers resistance to *Albugo candida* isolate AcEM2, whereas the clade 3 CHS3/CSA1 pair in Ws-2 is susceptible to AcEM2 (Parkes, 2020). This suggests that the ID acquisition is important for AcEM2 recognition by clade1 CHS3/CSA1. The clade 1 CHS3/CSA1 pair from Col-0 complements the AcEM2 susceptible phenotype of Ws-2, but the individual genes do not (Parkes, 2020), indicating that clade 1 Col-0 CSA1 or CHS3 cannot function with clade 3 Ws-2 CHS3 or CSA1. These results show that *CHS3* and *CSA1* have undergone co-evolution between clade1 on one hand and clades 2 and 3 on the other (Figure 7I).

Different configurations of divergence and co-evolution are observed across NLR pairs. Functional studies of the rice NLR Pias (Pias-2/Pias-1) pair, or the Pia (RGA5/RGA4) pair which are allelic to this pair, revealed that the executor NLRs Pias-1 or RGA4 are conserved, whereas only the sensor NLRs Pias-2 or RGA5 are highly divergent (Shimizu et al., 2021). Phylogenetic reconstruction of Pias/Pia pair showed that the sensor in each case has a higher level of divergence than the executor (Shimizu et al., 2021). In stark contrast, CSA1 and CHS3 tightly co-evolved to the extent that sensor and executor alleles from different clades within the same species cannot function or associate together, particularly in pairs where there is presence-absence of an ID in the sensor.

### Two single residue polymorphisms in CSA1 are required for CHS3/CSA1 complex function

We defined two polymorphic residues essential for CHS3/CSA1 pair function (Figures 7I). The G678/681 residue at the beginning of LRR domain is conserved in CSA1 alleles from all clades and is required for the function of CHS3/CSA1 pairs from all clades. CSA1 modeled onto the TNL RPP1 structure (Ma et al., 2020) revealed that this G residue is on the convex back surface of an LRR solenoid. It is possible that this CSA1 G residue is required to interact with the C terminus of CHS3 if the CHS3/CSA1 complex forms a tetramer like RPP1. In addition, we identified an aspartic acid (D543) residue conserved only in clade 2 and clade 3 CSA1 alleles. Structure modeling on the RPP1 showed that this CSA1 D residue is on the surface of a solenoid between the NB and the LRR. It is hard to predict its potential function. However, we found this residue is necessary to induce strong cell death and association with CHS3. Identification of these two residues indicates the evolution of both general and clade-specific residue functions (Figure 7I).

### The CHS3/CSA1 pair is regulated by two functional modes

Previous papers implicated a hypothetical NLR as a ‘Guard’ for HopB1-targeted BAK1 family members (Li et al., 2016; Wu et al., 2020). Consistent with this hypothesis, we found that the clade 1 CHS3/CSA1 Col-0 pair is necessary for *bak1 bkk1*-mediated auto-immune phenotype. We also demonstrated that BAK1 expression suppresses cell death caused by clade 2 or clade 3 CHS3/CSA1 pairs and that BAK1 associates with CSA1 from all clades. Recently, clade 1 CSA1_Col-0 was shown to be required for the auto-immune phenotype of *bak1 bir3*. It was identified as a direct interactor with BIR3 and as an indirect interactor with BAK1 (Schulze et al., 2021). We also found that BIRs associate with CSA1 and but only slightly suppress clade 2 and clade 3 CHS3/CSA1 pair-mediated cell death. Thus, our data are consistent with a model where clade 1 sensor CHS3 carrying an ID contributes to maintenance of the clade 1 CHS3/CSA1 pair in its inactive state when pathogen is absent, while both clade1-type and clade 2 / clade 3-type CHS3/CSA1 pairs ‘Guard’ various components of the BAK1-BIR complex (Figure 7I). In this model, association with BAK1-BIR holds the NLR pair in the inactive state until a pathogen infection alters the confirmation of the Guardee by cleaving BAK1 and its paralogs.

The paired TNL CHS1/SOC3 in Arabidopsis, which does not have an ID, triggers cell death when over-expressed and a PTI-regulating E3 ligase called SAUL1 is guarded by CHS1/SOC3 (Liang et al., 2019, 2020; Figure S2J). Phosphorylation of Thr1214 within the WRKY domain is required to keep RRS1-R in the autoinhibited state and dephosphorylation constitutively activates RPS4, leading to cell death and defense, so there might be another component regulating RRS1-R phosphorylation to maintain the complex in an inactive state in the absence of pathogen (Guo et al., 2020). Additionally, the sensor NLRs of rice Pik pairs, Pias pair and the *Brassica napus Bn*RPR2/*Bn*RPR1 pair are required for strong cell death induction of these pairs. Therefore, the previously defined functional mode of paired NLR regulation, where the sensor is an effector-activated negative regulator of an otherwise autonomous executor, as suggested for RRS1/RRS4 and RGA5/RGA4, may not be as prevalent as assumed. We predict that the “Guardee” negative regulatory mode for paired NLRs is common, and at least for the CHS3/CSA1 pair, ancestral (Figure S7I).

More pairs in Arabidopsis lack IDs than contain them (Van de Weyer et al., 2019). Our study therefore presents a novel case where we identify functionality for both pre-ID integration and post-ID integration NLR alleles that share a common ancestor. Furthermore, after ID integration, pre-ID integration functionality is maintained. However, there is a price for this maintenance in that the clade 1 alleles, with the ID, no longer function with clade 2 and clade 3 alleles that lack it. We hypothesize that the addition of the ID to clade 1 sensor CHS3 results in some evolutionary benefit, likely the recognition of an as yet unknown effector from *Albugo candida* isolate AcEM2. This mechanistic expansion likely comes with a cost under certain prevailing conditions, resulting in balancing selection at the CHS3/CSA1 locus.

## Acknowledgements

We thank Sarah Grant, Farid El-Kasmi and Marc T. Nishimura for excellent suggestions and comments, and Jia Li for providing the *bak1-3 bkk1-1* seeds. Supported by National Science Foundation (Grant IOS-1758400 to J.L.D.) and HHMI. J.L.D. is a Howard Hughes Medical Institute (HHMI) Investigator. N.H.K. was partially supported by Basic Science Research Program through the National Research Foundation of Korea Fellowship funded by the Ministry of Education (2014R1A6A3A03058629).

## Author contributions

Y.Y, O.J.F and J.L.D. conceptualized the project. J.L.D supervised the research. Y.Y led the experimentation. O.J.F. led the evolution and computation work. V.C provided truncated *CHS3_Col-0* constructs. All authors contributed to data analysis. Y.Y developed figures and wrote the main draft of the text. O.J.F and J.L.D. contributed significantly to that draft. N.H.K. helped to re-focus the figures and all authors contributed edits and comments to finalize the paper.

## METHODS

### Plant material and growth conditions

The several ecotypes of *Arabidopsis thaliana* were obtained from the *Arabidopsis* Biological Resource Center. *Arabidopsis* were grown in growth room at 21 °C/18 °C with 16h/8 h light/dark photoperiod on mixed soil. 3-5-week-old plants were used to extract genomic DNA or observe autoimmunity phenotype.

*Nicotiana benthamiana* (*N. benthamiana*) and *Nicotiana tabacum* (*N. tabacum*) were grown in growth room at 24 °C/20 °C under 16h/8 h light/dark photoperiod. 4-5-week-old plants were used to *Agrobacterium-* mediated transient expression for cell death phenotype, immunoblots and co-immunoprecipitation.

### Bacterial strains

*E. coli* Top 10 and *Agrobacterium tumefaciens* strain GV3101 were grown in LB media at 37 °C and 28 °C, respectively. Antibiotic concentrations used for *E. coli* and *Agrobacterium tumefaciens* were kanamycin 50 μg/mL, spectinomycin 50 μg/mL, gentamycin 25 μg/mL and rifampicin 100 μg/mL.

### Plasmid Constructions

Genomic fragments of CSA1, CHS3, BAK1, BKK1 and BIR were PCR amplified from *Arabidopsis thaliana* genomic DNA of indicated ecotypes (Figure S1B). The C-terminally HA-tagged, HF-tagged and V5-tagged constructs were generated using the Golden Gate assembly cloning procedure described previously and MYC-tagged constructs were generated using gateway assembly cloning procedure described previously (Ma et al., 2018). Briefly, for Golden Gate, the full-length genomic sequence was split into several short fragments for amplification, and PCR products, removing *Bsa I* sites without protein sequences change, were cloned into binary vector pICSL86922 with 35S promoter and TMV omega enhancer. For Gateway, binary vector pGWB617 was used carrying 35S promoter.

Site-directed mutants were generated by PCR mutagenesis. The full-length genomic sequence was split into two or more short fragments for amplification. Some primers carrying desired mutations were used to perform PCR. Then the PCR products were mixed and cloned into binary vector pICSL86922 using Golden Gate.

### *Agrobacterium*-mediated transient expressions

*Agrobacterium tumefaciens* strain GV3101 carrying indicated constructs were grown in liquid LB medium with relevant antibiotic for overnight in 28°C shaking incubator. Cell pellets were collected and re-suspended in infiltration buffer containing 10 mM MES (pH 5.6), 10 mM MgCl_2_, and 100 μM acetosyringone. Bacterial suspension was adjusted to OD_600_=0.5 in the final mix for infiltration. Leaves of 4-5-week-old *N. benthamiana* and *N. tabacum* were infiltrated with 1 mL needle-less syringe. Plants were put back in growth room at 24 °C/20 °C under 16h/8 h light/dark photoperiod after inoculation and cell death phenotypes were photographed at 4-5 dpi (days post infiltration). Leaves were harvested for immunoblots and Co-immunoprecipitation at 2 dpi.

### UV-light imaging of cell death phenotypes

4-5 days post infiltration leaves of *N. benthamiana* and *N. tabacum* were placed under UV lamps (B-100AP, UVP) and photographed using a digital camera with a yellow filter.

### Protein extraction and immunoblots

Leaves of *N. benthamiana* were harvested at 2 dpi and flash frozen in liquid nitrogen. Frozen samples were ground in liquid nitrogen and the powder was resuspended with an equal volume of extraction buffer (100 mM Tris-HCl pH 7.5, 1 mM EDTA pH 8.0, 150 mM NaCl, 1 % Triton X-100, 0.1% SDS, 10 mM dithiothreitol (DTT) and 1x Sigma plant protease inhibitor cocktail (Sigma-Aldrich)). Proteins were mixed with equal volume of 5x SDS-PAGE loading buffer and heated for 5 min at 95°C, then samples were centrifuged at 12000 rpm for 3 minutes. Samples were resolved in 8% SDS-PAGE gels and transferred to nitrocellulose membrane. Membranes were blocked with 5% milk dissolved in TBST and probed with primary anti-HA (Roche, 1:2000), anti-Flag (Sigma, 1:2000), Anti-V5 (Invitrogen, 1:5000) and Anti-MYC (Santa Cruz Biotechnology, 1:2000) and secondary HRP-conjugated anti-rat (Abcam, 1:10000) and anti-mouse (Santa Cruz Biotechnology, 1:10000) antibody.

### Co-immunoprecipitation (co-IP)

Leaves of *N. benthamiana* were harvested at 2 dpi and flash frozen in liquid nitrogen. Frozen samples were ground in liquid nitrogen and the powder was resuspended in 2 mL extraction buffer (50 mM HEPES pH 7.5, 150 mM NaCl, 10 mM EDTA pH 8.0, 0.5% Triton X-100, 5 mM DTT with 1× plant protease inhibitor mixture) and mixed well using vortex. Soluble supernatants were obtained by centrifugation twice at 10,400 × g for 5 min and 20,800 × g for 20 min at 4°C. Soluble supernatants were mixed with 25 μL of anti-HA or anti-MYC conjugated magnetic beads (Miltenyi Biotec) and incubated for 2 h with constant rotation at 4°C. The conjugated magnetic beads were captured using separation columns (Miltenyi Biotec) and were washed with washing buffer (50 mM HEPES pH 7.5, 150 mM NaCl, 10 mM EDTA pH 8.0, 0.2% Triton X- 100, 5 mM DTT with 1× plant protease inhibitor mixture) for three times. Proteins were eluted with 100 μL elution buffer (Miltenyi Biotec). Proteins were resolved in 8% SDS-PAGE gels described above.

### G and D residue prevalence in CSA1s and other TNLs

We investigated the incidence “G” and “D” in CSA1s and other TNLs across the plant phylogeny using publicly available protein sequence (http://ann-nblrrome.tuebingen.mpg.de/apollo/jbrowse/, https://github.com/weigelworld/pan-nlrome/). We aligned full-length protein sequence using MEGA7 and only kept some amino acids including the indicated “G” or “D” residue. Then we used the WebLogo website (https://weblogo.berkeley.edu/logo.cgi) to analyze residue prevalence.

### CRISPR/CAS9 construct design

The CRISPR/CAS9 constructs for targeting *CSA1* or *CHS3* were described previously (Wang et al., 2015). Briefly, *CSA1* and *CHS3* target sites were chosen using the CRISPR-PLANT and Cas-OFFinder websites (http://www.genome.arizona.edu/crispr/CRISPRsearch.html; http://www.rgenome.net/cas-offinder/). We designed two target sites to result in fragment deletions in *CSA1* and *CHS3*, respectively. The two gRNAs expression CRISPR cassette was generated by PCR amplification using pCBC-DT1T2 as template (CSA1-BsF 5’ - ATATATGGTCTCGATTGGGGGATAAGTTCAGGGAGCGTT - 3’, CSA1-F0 5’ - TGGGGGATAA GTTCAGGGAGCGTTTTAGAGCTAGAAATAGC - 3’, CSA1-R0 5’ - AACTGCAAGAATTTCGTGGCCACAATCTCTTAGTCGACTCTAC - 3’, CSA1-BsR 5’ - ATTATTGGTCTCGAAACTGCAAGAATTTCGTGGCCAC; CHS3-BsF 5’ - ATATATGGTCTCGATTGTTGCTTTACAAGCAACATAGTT - 3’, CHS3-F0 5’ – TGTTGC TTTACAAGCAACATAGTTTTAGAGCTAGAAATAGC - 3’, CHS3-R0 5’- AACTTAGCTCTCGGCCGTAAATCAATCTCTTAGTCGACTCTAC - 3’, CHS3-BsR 5’ – ATTATTGGTCTCGAAACTTAGCTCTCGGCCGTAA ATC - 3’). The PCR products were digested with *Bsa I* and cloned into binary vector pHEE401E using Golden Gate (Gao et al., 2013). These two recombinant constructs were transformed into *bak1-3 bkk1-1* plants by Agrobacterium-mediated floral-dip transformation (Clough & Bent, 1998), respectively. Genotyping and sanger sequencing were performed to identify transgenic plants with fragments deletion in *CSA1* and *CHS3*.

### Gene expression analysis

Total RNAs were isolated using the RNeasy Plant Mini Kit (Qiagen). RNase-Free DNase Set (Qiagen) was used to remove DNA. Complementary DNA was synthesized from 500 ng of total RNA using random primers (Thermofisher) and SuperScript™ III Reverse Transcriptase (Thermofisher). Real-time PCR was performed on the Applied Biosystems ViiA 7 using Power SYBR Green PCR Master Mix (Thermo Fisher Scientific). The level of *ACTIN7* (*ACT7*) expression was used as the internal control to normalize the *PR1* expression values. Biological replicates represented at least two independent experiments. Three technical replicates were performed for each experiment.

## Supplementary Figures

**Figure S1.**
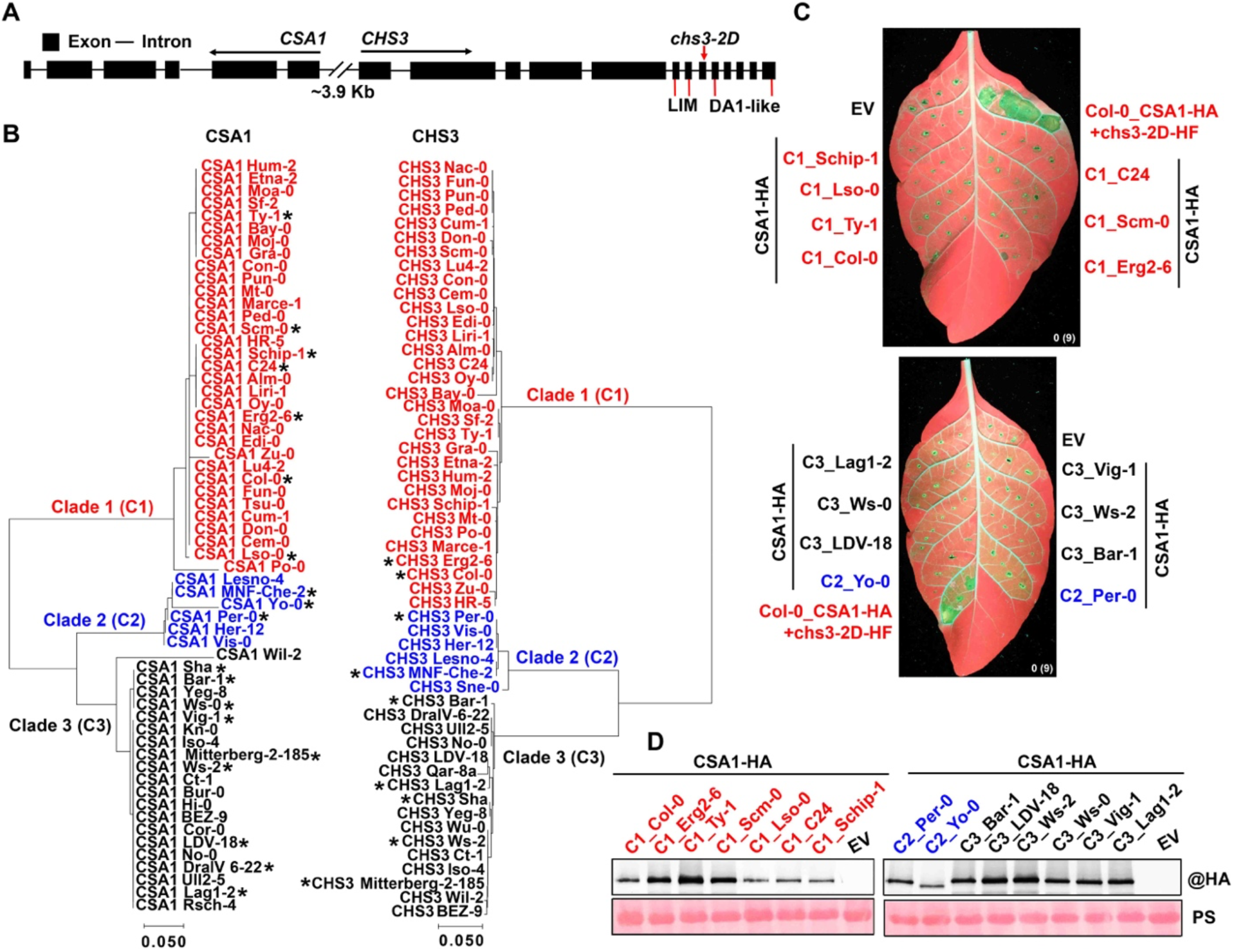
CSA1 expression alone cannot trigger evident cell death. (A) Schematic description of the *CSA1* and *CHS3* locus showing the position of the *chs3-2D* mutation and integrated domains (IDs) (LIM and DA1-like). (B) Neighbor-joining phylogenetic analysis and alignment of CSA1 (left) and CHS3 (right) based on the full-length amino acid sequence. CSA1 and CHS3 are divided into three clades (clade 1 or C1, clade 2 or C2 and clade 3 or C3). Clade 1 accessions and proteins are in red, clade 2 are in blue and clade 3 are in black. The scale bar indicates substitution per site. Asterisks mark all the accessions from which we cloned CSA1 or CHS3. (C and D) In planta (*N. tabacum*) phenotypes of CSA1 expression alone (C) and protein accumulation (D). Co-infiltration of Col-0 CSA1-HA with chs3-2D-HF is a positive control. Images were photographed at 4-5 dpi. EV: empty vector. The numbers represent the numbers of leaves with cell death out of the total number of leaves infiltrated. PS: ponceau stain indicates protein loading.

**Figure S2.**
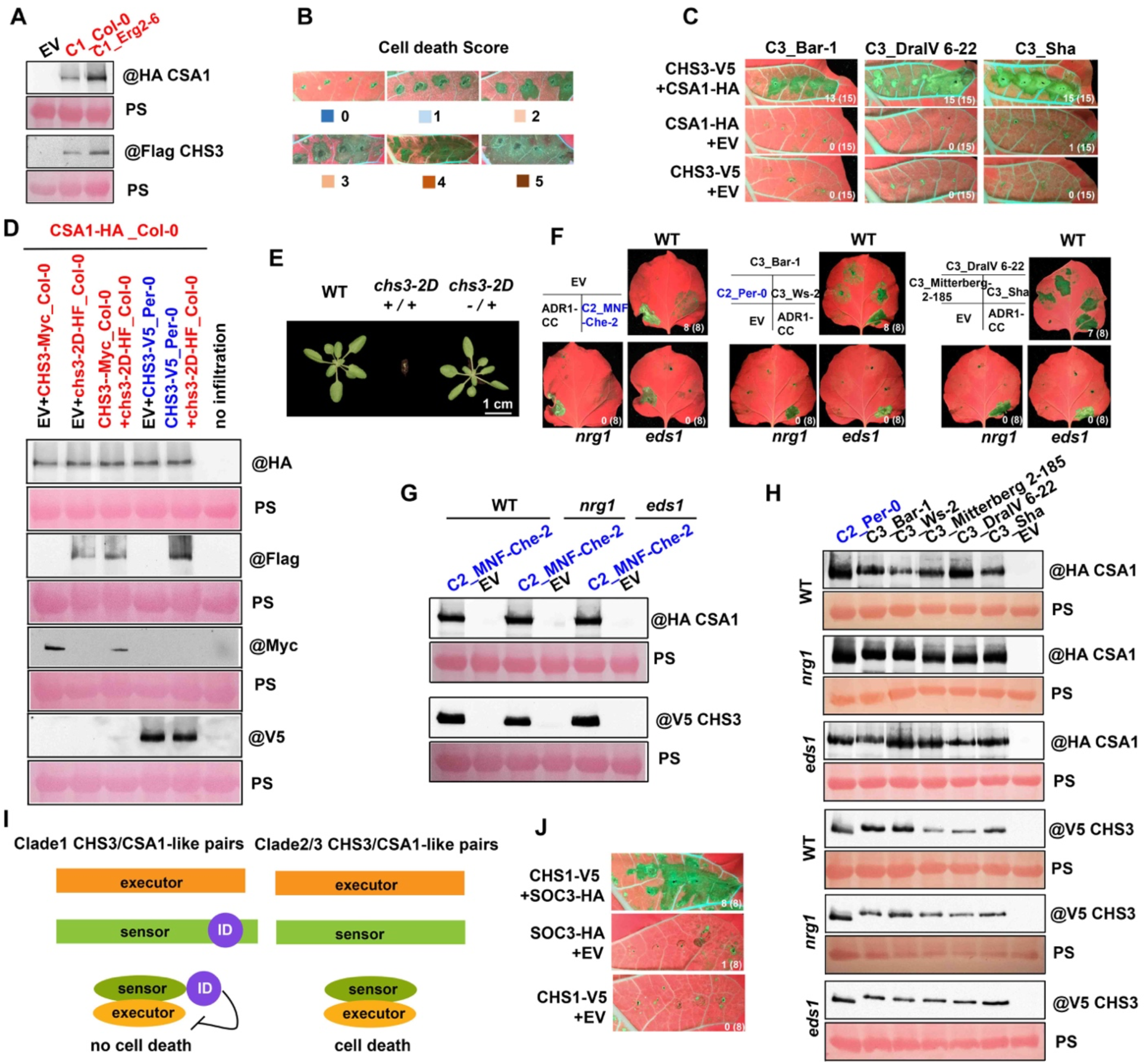
Wild-type CHS3/CSA1 pairs from clade 2 and clade 3 trigger EDS1- and NRG1-dependent cell death. (A) Immunoblot shows protein accumulation of CSA1 fused with HA tag and CHS3 fused with HF (6⊗His-3⊗Flag) tag from clade 1 Col-0 or Erg2-6 at 2 dpi in *N. benthamiana*. (B) Representative image of cell death score used throughout to determine the extent of cell death. (C) Clade 3 CHS3/CSA1 pairs trigger cell death in *N. tabacum*. Images were photographed at 4-5 dpi. EV: empty vector. The numbers represent the numbers of leaves displaying cell death out of the total number of leaves infiltrated. (D) Immunoblots confirm CSA1 and CHS3 proteins accumulation. EV: empty vector. PS: ponceau stain indicates protein loading. Clade 1 CSA1 and CHS3 are in red, clade 2 are in blue (E) Phenotypic analysis. Seedlings of indicated genotypes were grown in the 21°C daytime and 18°C night under long day (16h light/8h dark) condition. (F) Transient expression of clade 2 ad clade 3 matched CHS3/CSA1 pairs in either WT or *nrg1* or *eds1* mutant *N. benthamiana*. Images were photographed at 4-5 dpi. Infiltration of ADR1-CC and EV (empty vector) is a positive and negative control, respectively. Accessions of clade 2 show in blue and clade 3 show in black. The numbers indicate the numbers of leaves showing cell death out of the total number of leaves infiltrated. (G and H) Western blots confirm protein accumulation of CHS3/CSA1 pairs in indicated *Nb* genotypes. EV: empty vector. Accessions of clade 2 show in blue and clade 3 show in black. (I) Schematic representation of the proposed cell death induction by two different types of NLR pairs. (J) In planta (*N. tabacum*) phenotypes of paired TNL CHS1/SOC3. Images were photographed at 4-5 dpi. EV: empty vector. The numbers indicate the numbers of leaves showing cell death out of the total number of leaves infiltrated.

**Figure S3.**
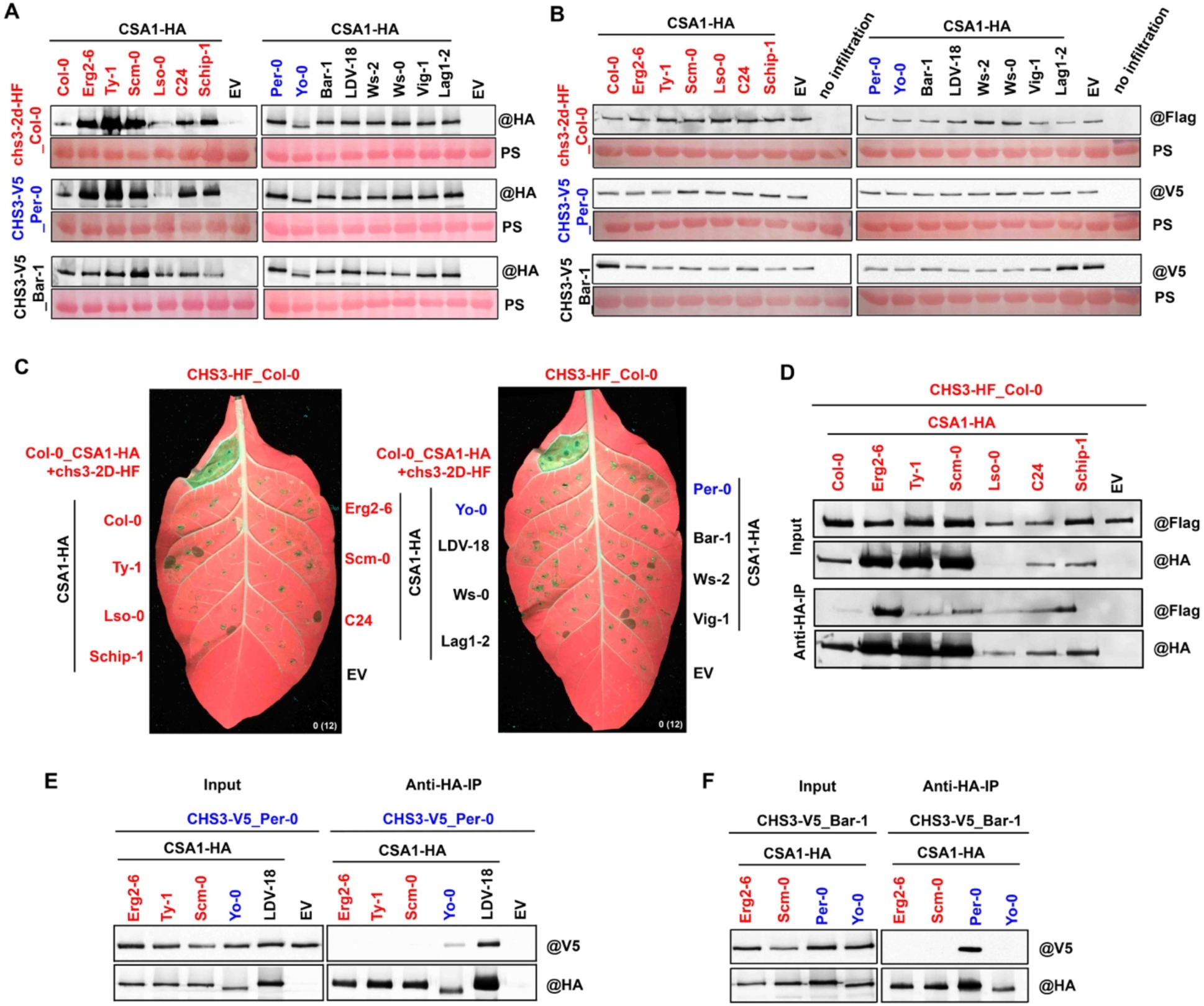
Intra- and inter-clade combinations of clade 1 CHS3_Col-0 with CSA1 do not trigger cell death and clade 1 CSA1 cannot associate with clade 2 CHS3_Per-0 or clade 3 CHS3_Bar-1. (A and B) Immunoblots show protein accumulation of CSA1 (A) and CHS3 (B). EV: empty vector. PS: ponceau stain indicates protein loading. Clade 1 accessions and proteins are in red, clade 2 are in blue and clade 3 are in black, respectively. (C) In planta (*N. tabacum*) phenotypes. Images were photographed at 4-5 dpi. Co-infiltration of Col-0 CSA1-HA with chs3-2D-HF is a positive control. Clade 1 accessions and proteins are in red, clade 2 are in blue and clade 3 are in black, respectively. (D) Co-IP assays showing all clade 1 CSA1s we tested associate with wild-type CHS3_Col-0, even though they do not induce cell death. Clade 1 CSA1 and CHS3 are in red. (E and F) Co-IP assays reveal that V5-tagged clade 2 CHS3_Per-0 and clade 3 CHS3_Ws-2 do not associate with HA-tagged clade 1 CSA1. EV: empty vector. Clade 1 accessions and proteins are in red, clade 2 are in blue and clade 3 are in black.

**Figure S4.**
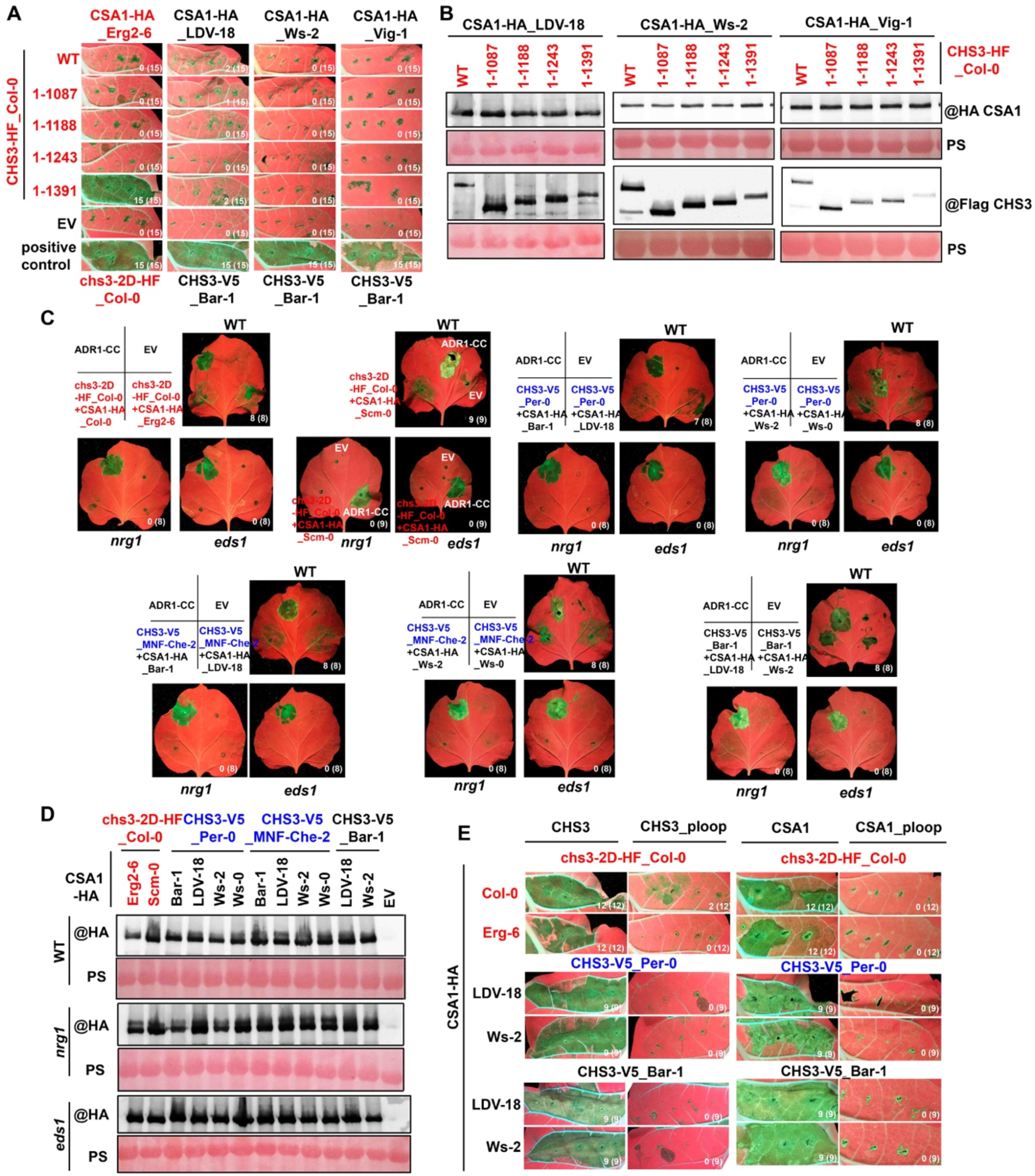
The combination of the DA1-like ID truncation of clade 1 CHS3 with clade 1 but not clade 3 CSA1 leads to cell death and all combinations trigger EDS1- and NRG1-dependent cell death that requires intact CHS3 and CSA1 p-loops. (A) Transient co-expression of clade 1 HF-tagged full-length or truncated CHS3_Col-0 with HA-tagged CSA1. Images were photographed at 4-5 dpi. The numbers represent the numbers of leaves displaying cell death out of the total number of leaves infiltrated. (B) Immunoblots confirm protein accumulation. Constructs are blotted with either anti-HA (CSA1) or anti-Flag (CHS3). PS: ponceau stain indicates protein loading. (C) UV-image of *N*.*benthamiana* leaves of indicated genotypes WT, *nrg1* and *eds1* expressing different CHS3/CSA1 combinations. ADR1-CC is a positive control and EV is empty vector negative control. Images were photographed at 4-5 dpi. Clade 1 accessions and proteins are in red, clade 2 are in blue and clade 3 are in black, respectively. The numbers represent the numbers of leaves with cell death out of the total number of leaves infiltrated. (D) Immunoblots show protein accumulation of HA-tagged CSA1. Samples were harvested at 2 dpi. EV: empty vector. PS: ponceau stain indicates protein loading. Clade 1 accessions and proteins are in red, clade 2 are in blue and clade 3 are in black, respectively. (E) Cell death assays reveal both intact p-loops of CSA1 and CHS3 are essential for intra- and inter-clade CHS3/CSA1 combinations to trigger cell death in *N. tabacum*. Images were photographed at 4-5 dpi. Clade 1 accessions and proteins are in red, clade 2 are in blue and clade 3 are in black, respectively.

**Figure S5.**
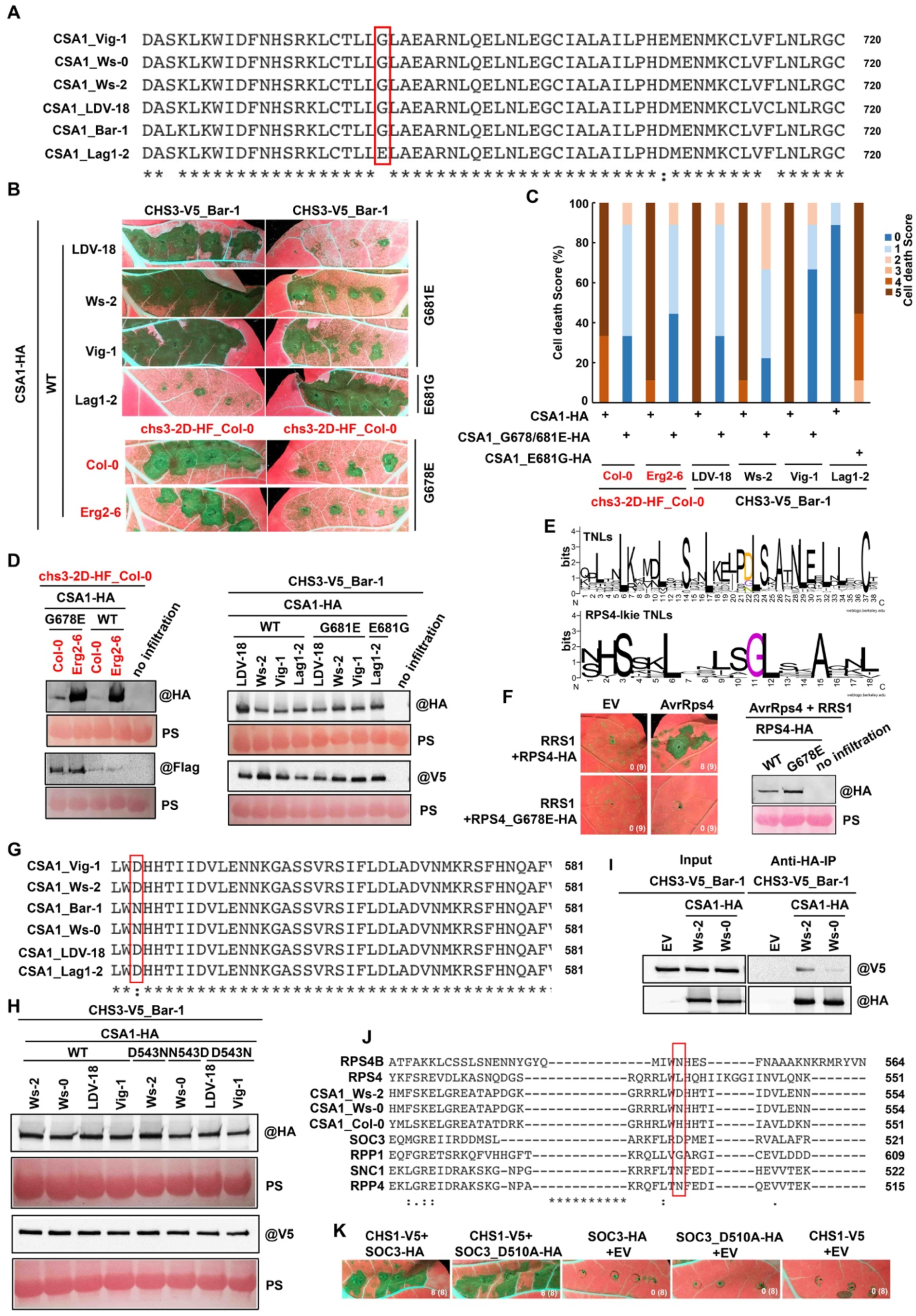
Two single residue polymorphisms in CSA1 are required for CHS3/CSA1 complex function. (A) Amino acid sequence alignment of indicated clade 3 CSA1s showing the conservation of G681 in CSA1s from clade 3, except CSA1_Lag1-2. Red box indicates the position of G residue. (B) In planta phenotypes. Images were photographed at 4-5 dpi. Clade 1 accessions and proteins show in red and clade 3 show in black, respectively. (C) Percentage representations of cell death score in (B). Stacked bars are color-coded showing the proportions (in percentage) of each cell death score (0–5). 9 leaves were scored for each stacked bar. (D) Protein accumulation of CHS3, CSA1 and mutant of CSA1. PS: ponceau stain indicates protein loading. (E) A sequence logo showing conserved G residue in LRR domain of RPS4-like TNL. (F) In planta phenotypes and protein accumulation. G residue is required for RRS1/RPS4 pair to recognize AvrRps4 (delivered via Agrobacterium) and trigger cell death in *N. benthamiana*. (G) Amino acid sequence alignment of indicated clade 3 CSA1s. Red box shows the position of D543. (H) Immunoblots confirm protein accumulation. PS: ponceau stain indicates protein loading. (I) Co-IP assays reveal that the association of clade 3 V5-tagged CHS3_Bar-1 with clade 3 HA-tagged CSA1_Ws-2 is stronger than with clade 3 HA-tagged CSA1_Ws-0. Constructs are blotted with either anti-HA (CSA1) or anti-V5 (CHS3). (J) Protein sequence alignment of indicated TNLs reveals the conserved D residue in SOC3. (K) Cell death assays showing the conserved D residue in SOC3 is not required for CHS1/SOC3-mediated cell death. Images were photographed at 4-5 dpi. The numbers represent the numbers of leaves displaying cell death out of the total number of leaves infiltrated.

**Figure S6.**
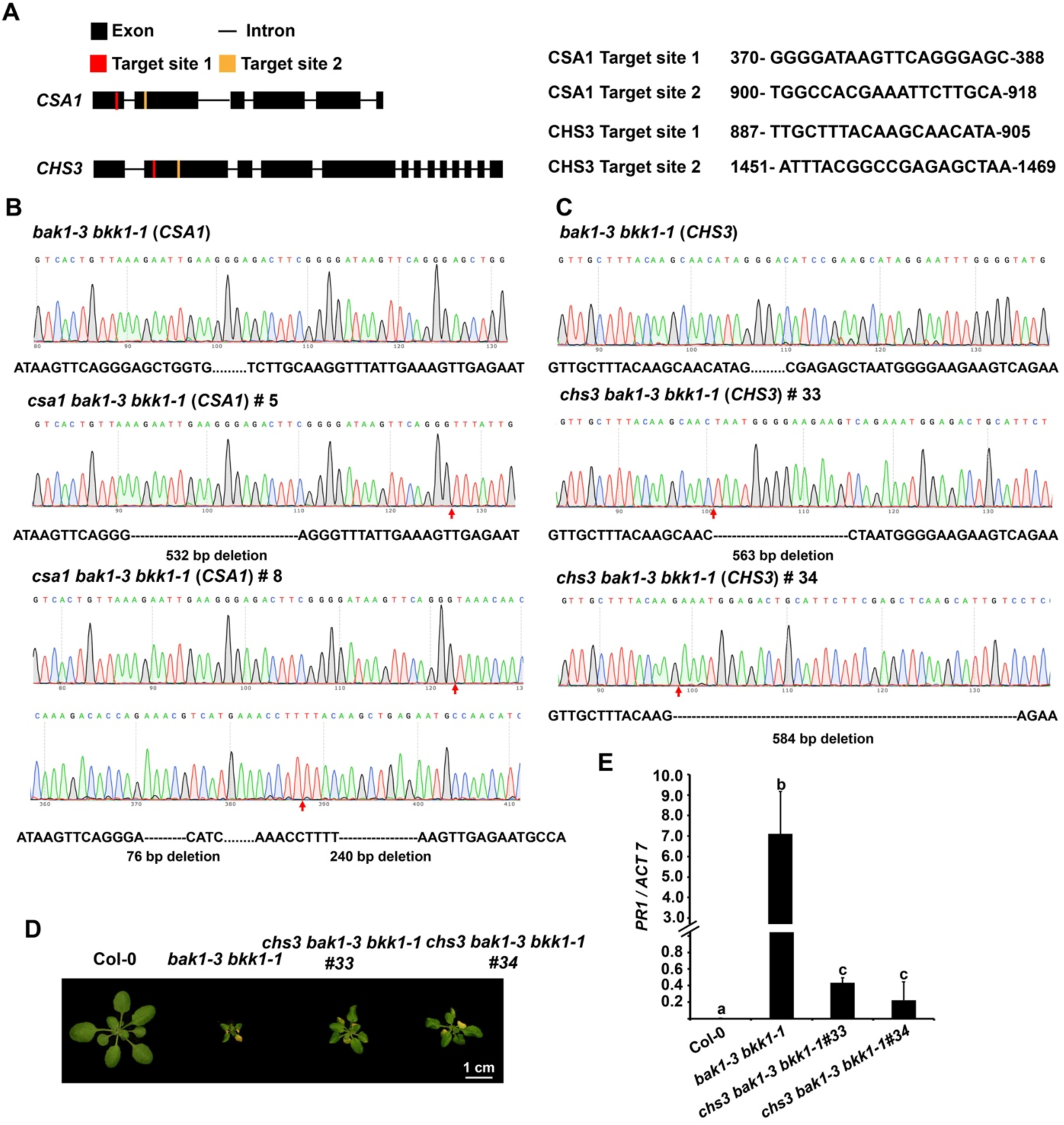
Identification of triple mutants of *csa1 bak1-3 bkk1-1* and *chs3 bak1-3 bkk1-1*. (A) Schematic description of the CRISPR/CAS9 design for deleting CSA1 and CHS3 (left) and targeting position (right). Boxes indicate exons and lines indicate introns. Red boxes represent target site 1 and yellow boxes represent target site 2. (B and C) Sanger sequencing chromatograph and genomic DNA sequence comparison between wild-type CSA1 (B)/CHS3 (C) and mutated CSA1 (B)/CHS3 (C). (D and E) Morphological phenotypes (D) and *PR1* gene expression (E) of indicated genotypes are shown. *chs3* mutation partially suppresses *bak1 bkk1*-mediated auto-immune phenotype and increased *PR1* gene expression. Plants were grown in 21 °C/18 °C under LD (16 h light/8 h dark) condition. Images were photographed at 3-5 weeks. Error bars represent two biological repeats (three technical replicates were performed for each biological repeat). The different letters “a-c” indicate statistically significant differences determined by one-way ANOVA for multiple pairwise comparisons and Tukey’s least significant difference test. P<0.05.

**Figure S7.**
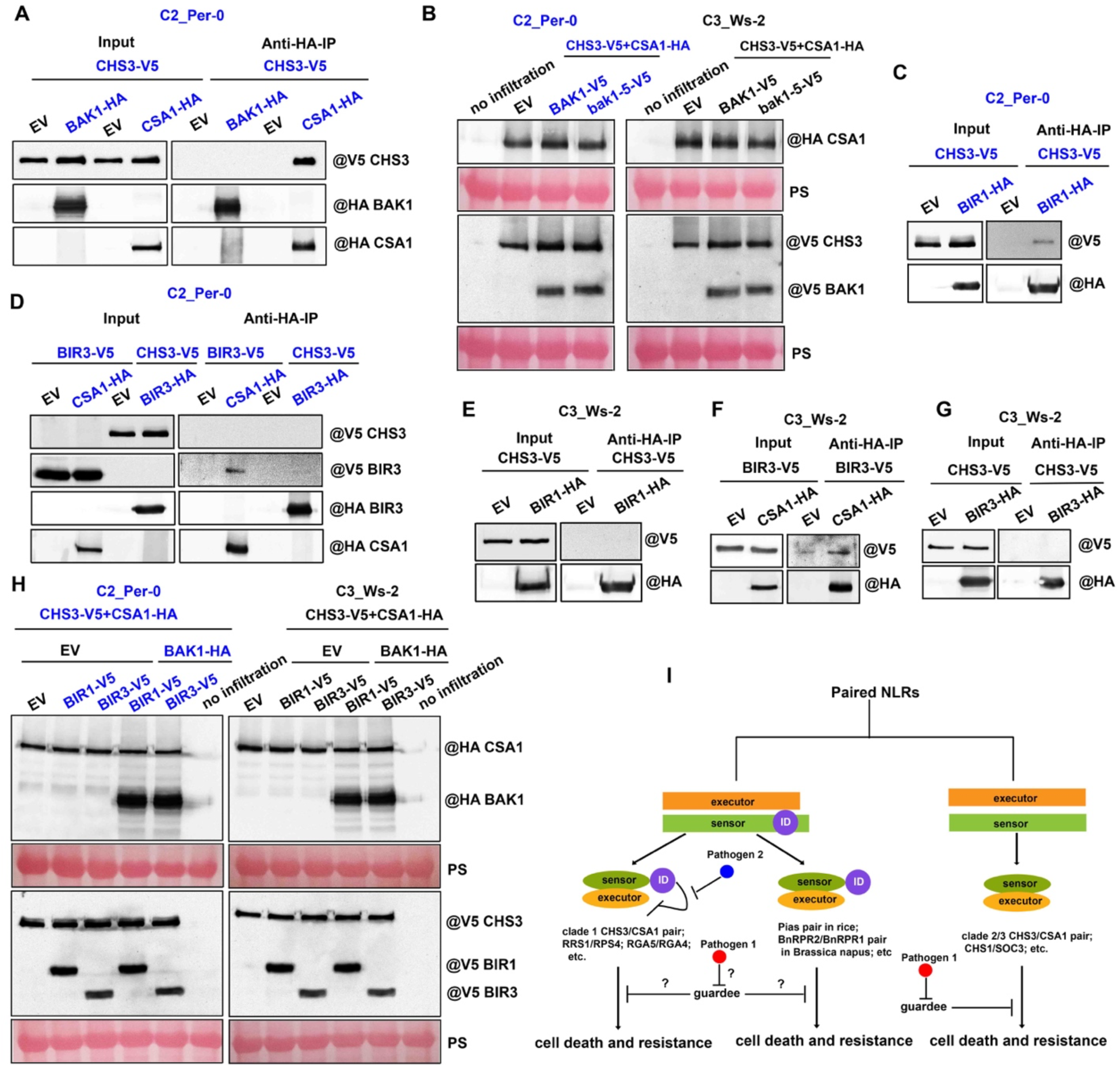
BAK1 and BIR differentially associate with CHS3/CSA1 pairs. (A) Co-IP assays show no association between BAK1 and clade 2 CHS3_Per-0. Total proteins were extracted from *N. benthamiana* at 2 dpi. Total proteins (input) or immunoprecipitation products were probed in immunoblots with antibodies to V5 or HA. Co-expression of clade 2 CSA1_Per-0 and CHS3_Per-0 is a positive control. (B) Western blots confirm protein accumulation of HA-tagged CSA1 and V5-tagged CHS3 and BAK1. PS: ponceau stain indicates protein loading. (C-G) Co-IP assays assess the association of BIR with CSA1 and CHS3 of clade 2 and clade 3. BIR1 (C and E) and BIR3 (D, F and G) enable associate with clade 2 CSA1_Per-0 and clade 3 CSA1_Ws-2, but weak or no association with CHS3s. Total proteins were extracted from *N. benthamiana* at 2 dpi. (H) Immunoblots confirm protein accumulation of HA-tagged CSA1 and BAK1 and V5-tagged CHS3 and BIRs. PS: ponceau stain indicates protein loading. (I) Hypothetical model for paired NLR regulation. A subset of paired NLRs have sensors that contain IDs and act negative regulators. These might have two ways to maintain the inactive state in the absence of pathogen, meanwhile there are two ways to recognize pathogens and activate the complex. One mediated by sensors, and the other mediated via negative regulation by a guardee, such as clade 1 CHS3/CSA1 pairs. For other paired NLRs, even with ID in the sensors, the wild-type pair triggers cell death, suggesting that the “guardee” might be indispensable to maintain their resting state, such as clade 2/3 CHS3/CSA1 pair.

## Notes

### Competing Interest Statement

The authors have declared no competing interest.

## References Cited

Adachi, H., Derevnina, L., and Kamoun, S. (2019). NLR singletons, pairs, and networks: evolution, assembly, and regulation of the intracellular immunoreceptor circuitry of plants. Curr. Opin. Plant Biol. 50, 121–131.

Bayless, A.M., Nishimura, M.T. (2020) Reinventing the wheel with a synthetic plant inflammasome. Proc. Natl. Acad. Sci. USA. 117, 20357–20359.

Bi, D., Johnson, K.C., Zhu, Z., Huang, Y., Chen, F., Zhang, Y., and Li, X. (2011). Mutations in an atypical TIR-NB-LRR-LIM resistance protein confer autoimmunity. Front. Plant Sci. 2, 71.

Bi, G., Su, M., Li, N., Liang, Y., Dang, S., Xu, J., Hu, M., Wang, J., Zou, M., Deng, Y., et al. (2021). The ZAR1 resistosome is a calcium-permeable channel triggering plant immune signaling. Cell 184, 3528–3541.

Boller, T., and Felix, G. (2009). A renaissance of elicitors: perception of microbe-associated molecular patterns and danger signals by pattern-recognition receptors. Annu. Rev. Plant Biol. 60, 379–406.

Castel, B., Ngou, P.-M., Cevik, V., Redkar, A., Kim, D.-S., Yang, Y., Ding, P., and Jones, J.D. (2019). Diverse NLR immune receptors activate defence via the RPW8-NLR NRG1. New Phytol. 222, 966–980.

Cesari, S., Bernoux, M., Moncuquet, P., Kroj, T., and Dodds, P.N. (2014a). A novel conserved mechanism for plant NLR protein pairs: the “integrated decoy” hypothesis. Front. Plant Sci. 5, 606.

Césari, S., Kanzaki, H., Fujiwara, T., Bernoux, M., Chalvon, V., Kawano, Y., Shimamoto, K., Dodds, P., Terauchi, R., and Kroj, T. (2014b). The NB-LRR proteins RGA4 and RGA5 interact functionally and physically to confer disease resistance. EMBO J. 33, 1941–1959.

Dodds, P.N., and Rathjen, J.P. (2010). Plant immunity: towards an integrated view of plant-pathogen interactions. Nat. Rev. Genet. 11, 539–548.

Clark, R.M., Schweikert, G., Toomajian, C., Ossowski, S., Zeller, G., Shinn, P., Warthmann, N., Hu, T.T., Fu, G., Hinds, D.A., et al. (2007). Common sequence polymorphisms shaping genetic diversity in Arabidopsis thaliana. Science 317, 338–342.

Clough, S.J., Bent, A.F. (1998). Floral dip: a simplified method for Agrobacterium-mediated transformation of Arabidopsis thaliana. Plant J.16, 735–743.

Couto, D., and Zipfel, C. (2016). Regulation of pattern recognition receptor signalling in plants. Nat. Rev. Immunol. 16, 537–552.

Cui, H., Tsuda, K., and Parker, J.E. (2015). Effector-triggered immunity: from pathogen perception to robust defense. Annu. Rev. Plant Biol. 66, 487–511.

De la Concepcion, J.C., Benjumea, J.V., Bialas, A., Terauchi, R., Kamoun, S., and Banfield, M.J. (2021). Functional diversification gave rise to allelic specialization in a rice NLR immune receptor pair. Elife 10, e71662.

Dominguez-Ferreras, A., Kiss-Papp, M., Jehle, A.K., Felix, G., and Chinchilla, D. (2015). An overdose of the Arabidopsis coreceptor BRASSINOSTEROID INSENSITIVE1-ASSOCIATED RECEPTOR KINASE1 or its ectodomain causes autoimmunity in a SUPPRESSOR OF BIR1-1-dependent manner. Plant Physiol. 168, 1106–1121.

Faigón-Soverna, A., Harmon, F.G., Storani, L., Karayekov, E., Staneloni, R.J., Gassmann, W., Más, P., Casal, J.J., Kay, S.A., and Yanovsky, M.J. (2006). A constitutive shade-avoidance mutant implicates TIR-NBS-LRR proteins in Arabidopsis photomorphogenic development. Plant Cell 18, 2919–2928.

Gao, X., Yan, P., Shen, W., Li, X., Zhou, P., Li, Y. (2013). Modular construction of plasmids by parallel assembly of linear vector components. Anal Biochem. 437, 172–7.

Gao, Y., Wu, Y., Du, J., Zhan, Y., Sun, D., Zhao, J., Zhang, S., Li, J., and He, K. (2017). Both light-Induced SA accumulation and ETI mediators contribute to the cell death regulated by BAK1 and BKK1. Front. Plant Sci. 8, 622.

Guo, H., Ahn, H.-K., Sklenar, J., Huang, J., Ma, Y., Ding, P., Menke, F.L., and Jones, J.D.G. (2020). Phosphorylation-regulated activation of the Arabidopsis RRS1-R/RPS4 immune receptor complex reveals two distinct effector recognition mechanisms. Cell Host Microbe 27, 769–781.

Guo, H., Wang, S., and Jones, J.D.G. (2021). Autoactive Arabidopsis RPS4 alleles require partner protein RRS1-R. Plant Physiol. 185, 761–764.

He, K., Gou, X., Yuan, T., Lin, H., Asami, T., Yoshida, S., Russell, S.D., and Li, J. (2007). BAK1 and BKK1 regulate brassinosteroid-dependent growth and brassinosteroid-independent cell-death pathways. Curr. Biol. 17, 1109–1115.

He, K., Gou, X., Powell, R.A., Yang, H., Yuan, T., Guo, Z., and Li, J. (2008). Receptor-like protein kinases, BAK1 and BKK1, regulate a light-dependent cell-death control pathway. Plant Signal. Behav. 3, 813–815.

Huang, S., Jia, A., Song, W., Hessler, G., Meng, Y., Sun, Y., Xu, L., Laessle, H., Jirschitzka, J., Ma, S., et al. (2022). Identification and receptor mechanism of TIR-catalyzed small molecules in plant immunity. BioRxiv. doi: https://doi.org/10.1101/2022.04.01.486681.

Huh, S.U., Cevik, V., Ding, P., Duxbury, Z., Ma, Y., Tomlinson, L., Sarris, P.F., Jones, J.D. (2017). Protein-protein interactions in the RPS4/RRS1 immune receptor complex. PLoS Pathog. 13, e1006376.

Jacob, Pierre., Kim, N.H., Wu, F., EI-Kasmi, F., Chi, Yuan., Walton, W.G., Furzer, O.J., Lietzan, A.D., Sunil, S., Kempthorn, K., et al. (2021). Plant “helper” immune receptors are Ca^2+^-permeable nonselective cation channels. Science 373, 420–425.

Jones, J.D.G., and Dangl, J.L. (2006). The plant immune system. Nature 444, 323–329.

Jones, J.D.G., Vance, R.E., and Dangl, J.L. (2016). Intracellular innate immune surveillance devices in plants and animals. Science 354, aaf6395.

Jubic, L.M., Saile, S., Furzer, O.J., El Kasmi, F., Dangl, J.L. (2019). Help wanted: helper NLRs and plant immune responses. Curr Opin Plant Biol. 50, 82–94.

Kemmerling, B., Schwedt, A., Rodriguez, P., Mazzotta, S., Frank, M., Qamar, S.A., Mengiste, T., Betsuyaku, S., Parker, J.E., Müssig, C., et al. (2007). The BRI1-associated kinase 1, BAK1, has a brassinolide-independent role in plant cell-death control. Curr. Biol. 17, 1116–1122.

Kumar, S., Stecher, G., and Tamura, K. (2016). MEGA7: molecular Evolutionary genetics analysis version 7.0 for bigger datasets. Mol. Biol. Evol. 33,1870–1874.

Li, L., Kim, P., Yu, L., Cai, G., Chen, S., Alfano, J.R., Zhou, J-.M. (2016). Activation-dependent destruction of a co-receptor by a Pseudomonas syringae effector dampens plant immunity. Cell Host Microbe 20, 504–514.

Liang, W., Wersch, S.V., Tong, M., and Li, X. (2019). TIR-NB-LRR immune receptor SOC3 pairs with truncated TIR-NB protein CHS1 or TN2 to monitor the homeostasis of E3 ligase SAUL1. New Phytol. 221, 2054–2066.

Liang, W., Tong, M., and Li, X. (2020). SUSA2 is an F-box protein required for autoimmunity mediated by paired NLRs SOC3-CHS1 and SOC3-TN2. Nat. Commun. 11, 5190.

Macho, A.P., and Zipfel, C. (2014). Plant PRRs and the activation of innate immune signaling. Mol. Cell 54, 263–272.

Ma, Y., Guo, H., Hu, L., Martinez, P.P., Moschou, P.N., Cevik, V., Ding, P., Duxbury, Z., Sarris, P.F., and Jones, J.D. (2018). Distinct modes of derepression of an Arabidopsis immune receptor complex by two different bacterial effectors. Proc. Natl. Acad. Sci. USA 115, 10218–10227.

Ma, S., Lapin, D., Liu, L., Sun, Y., Song, W., Zhang, X., Logemann, E., Yu, D., Wang, J., Jirschitzka, J., et al. (2020). Direct pathogen-induced assembly of an NLR immune receptor complex to form a holoenzyme. Science 370, eabe3069.

Martin, R., Qi, T., Zhang, H., Liu, F., King, M., Toth, C., Nogales, E., and Staskawicz, B.J. (2020). Structure of the activated ROQ1 resistosome directly recognizing the pathogen effector XopQ. Science 370, eabd9993.

Meyers, B.C., Dickerman, A.W., Michelmore, R.W., Sivaramakrishnan, S., Sobral, B.W., Young, N.D. (1999). Plant disease resistance genes encode members of an ancient and diverse protein family within the nucleotide-binding superfamily. Plant J. 20, 317–32.

Mermigka, G., Amartolou, A., Mentzelopoulou, A., Astropekaki, N., Sarris, P.-F. (2021). Assassination Tango: An NLR/NLR-ID immune receptors pair of rapeseed co-operates inside the nucleus to activate cell death. BioRxiv. https://www.biorxiv.org/content/10.1101/2021.10.29.466428v1.

Mukhtar, M.S., Carvunis, A.R., Dreze, M., Epple, P., Steinbrenner, J., Moore, J., Tasan, M., Galli, M., Hao, T., Nishimura, M.T., et al. (2011). Independently evolved virulence effectors converge onto hubs in a plant immune system network. Science 333, 596–601.

Ngou, B.P., Ahn, H.-K., Ding, P., and Jones, J.D. (2021). Mutual potentiation of plant immunity by cell-surface and intracellular receptors. Nature 592, 110–115.

Parkes, T., (2020). From Recognition to Susceptibility: Functional characterization of Plant-specific LIM-domain containing proteins in plant-microbe interactions. PhD thesis, University of Bath, Bath, UK

Pruitt, R.N., Locci, F., Wanke, F., Zhang, L., Saile, S.C., Joe, A., Karelina, D., Hua, C., Fröhlich, K., Wan, W.L., et al. (2021). The EDS1-PAD4-ADR1 node mediates Arabidopsis pattern-triggered immunity. Nature 598, 495–499.

Saraste, M., Sibbald, P.R., Wittinghofer, A. (1990) The P-loop a common motif in ATP-and GTP-binding proteins. Tr. Biochem. Sci. 15, 430–434.

Saucet, S.B., Ma, Y., Sarris, P.F., Furzer, O.J., Sohn, K.H., and Jones, J.D. (2015). Two linked pairs of Arabidopsis TNL resistance genes independently confer recognition of bacterial effector AvrRps4. Nat. Commun. 6, 6338.

Sarris, P.F., Duxbury, Z., Huh, S.U., Ma, Y., Segonzac, C., Sklenar, J., Derbyshire, P., Cevik, V., Rallapalli, G., Saucet, S.B., et al. (2015). A Plant immune receptor detects pathogen effectors that target WRKY transcription factors. Cell 161, 1089–1100.

Schwessinger, B., Boux, M., Kadota, Y., Ntoukakis, V., Sklenar, J., Jones, A., and Zipfel, C. (2011). Phosphorylation-dependent differential regulation of plant growth, cell death, and innate immunity by the regulatory receptor-like kinase BAK1. PLoS Genet. 7, e1002046.

Schulze, S., Yu, L., Ehinger, A., Kolb, D., Saile, S.C., Stahl, M., Franz-Wachtel, M., Li, L., Kasmi, F.E., Cevik, V., et al. (2021). The TIR-NBS-LRR protein CSA1 is required for autoimmune cell death in Arabidopsis pattern recognition co-receptor bak1 and bir3 mutants. BioRxiv. https://doi.org/10.1101/2021.04.11.438637.

Shimizu, M., Hirabuchi, A., Sugihara, Y., Abe, A., Takeda, T., Kobayashi, M., Hiraka, Y., Kanzaki, E., Oikawa, K., Saitoh, H., et al. (2021). A genetically linked pair of NLR immune receptors show contrasting patterns of evolution. BioRxiv. https://doi.org/10.1101/2021.09.01.458560.

Tajima, F. (1989). Statistical method for testing the neutral mutation hypothesis by DNA polymorphism. Genetics 123, 585–595.

Tian, H., Wu, Z., Chen, S., Ao, K., Huang, W., Yaghmaiean, H., Sun, T., Xu, F., Zhang, Y., Wang, S., et al. (2021). Activation of TIR signalling boosts pattern-triggered immunity. Nature 598, 500–503.

Van der Hoorn, R.A., and Kamoun, S. (2008). From guard to decoy: a new model for perception of plant pathogen effectors. Plant Cell 20, 2009–2017.

Van de Weyer, A.-L., Monteiro, F., Furzer, O.J., Nishimura, M.T., Cevik, V., Witek, K., Jones, J.D., Dangl, J.L., Weigel, D., and Bemm, F. (2019). A species-wide inventory of NLR genes and alleles in Arabidopsis thaliana. Cell 178, 1260–1272.

Walker, J.E., Saraste, M., Runswick, M.J., Gay, N.J. (1982) Distantly related sequences in the α-and β-subunits of ATP synthetase, myosin, kinases and other ATP-requiring enzymes and a common nucleotide binding fold. EMBO J. 1, 945–951.

Wang, Z.P., Xing, H.L., Dong, L., Zhang, H.Y., Han, C.Y., Wang, X.C., Chen, Q.J. (2015). Egg cell-specific promoter-controlled CRISPR/Cas9 efficiently generates homozygous mutants for multiple target genes in Arabidopsis in a single generation. Genome Biol. 16, 144.

Wang, J., Hu, M., Wang, J., Qi, J., Han, Z., Wang, G., Qi, Y., Wang, H.-W., Zhou, J.-M., and Chai, J. (2019a). Reconstitution and structure of a plant NLR resistosome conferring immunity. Science 364, eaav5870.

Wang, J., Wang, J., Hu, M., Wu, S., Qi, J., Wang, G., Han, Z., Qi, Y., Gao, N., Wang, H.-W., et al. (2019b). Ligand-triggered allosteric ADP release primes a plant NLR complex. Science 364, eaav5868.

Wan, L., Essuman, K., Anderson, R.G., Sasaki, Y., Monteiro, F., Chung, E.-H., Nishimura, E.O., DiAntonio, A., Milbrandt, J., Dangl, J.L., et al. (2019). TIR domains of plant immune receptors are NAD^+^-cleaving enzymes that promote cell death. Science 365, 799–803.

Wani, S.H., Anand, S., Singh, B., Bohra, A., Joshi, R. (2021). WRKY transcription factors and plant defense responses: latest discoveries and future prospects. Plant Cell Rep. 40, 1071–1085.

Weßling, R., Epple, P., Altmann, S., He, Y., Yang, L., Henz, S.R., McDonald, N., Wiley, K., Bader, K.C., Gläßer, C., et al. (2014). Convergent targeting of a common host protein-network by pathogen effectors from three kingdoms of life. Cell Host Microbe 16, 364–375.

Williams, S.J., Sohn, K.H., Wan, L., Bernoux, M., Sarris, P.F., Segonzac, C., Ve, T., Ma, Y., Saucet, S.B., Ericsson, D.J., et al. (2014) Structural basis for assembly and function of a heterodimeric plant immune receptor. Science 34, 299–303

Wirthmueller, L., Zhang, Y., Jones, J.D., and Parker, J.E. (2007). Nuclear accumulation of the Arabidopsis immune receptor RPS4 is necessary for triggering EDS1-dependent defense. Curr. Biol. 17, 2023–2029.

Wu, C.-H., Derevnina, L., and Kamoun, S. (2018). Receptor networks underpin plant immunity. Science 360, 1300–1301.

Wu, Z., Li, M., Dong, O.X., Xia, S., Liang, W., Bao, Y., Wasteneys, G., and Li, X. (2019). Differential regulation of TNL-mediated immune signaling by redundant helper CNLs. New Phytol. 222, 938–953.

Wu, Y., Gao, Y., Zhan, Y., Kui, H., Liu, H., Yang, L., Kemmerling, B., Zhou, J.M., He, Kai., Li, J. (2020). Loss of the common immune coreceptor BAK1 leads to NLR-dependent cell death. Proc. Natl. Acad. Sci. USA 117, 27044–27053

Xu, F., Zhu, C., Cevik, V., Johnson, K., Liu, Y., Sohn, K., Jones, J.D., Holub, E.B., and Li, X. (2015). Autoimmunity conferred by chs3-2D relies on CSA1, its adjacent TNL-encoding neighbour. Sci. Rep. 5, 8792.

Yuan, M., Jiang, Z., Bi, G., Nomura, K., Liu, M., Wang, Y., Cai, B., Zhou, J.-M., He, S.Y., and Xin, X.-F. (2021). Pattern-recognition receptors are required for NLR-mediated plant immunity. Nature 592, 105–109.

Yu, D., Song, W., Tan E.Y.J., Liu, L., Cao, Y., Jirschitzka, J., Li, E., Logemann, E., Xu, C., Huang, S., et al. (2021). TIR domains of plant immune receptors are 2′,3′-cAMP/cGMP synthetases mediating cell death. BioRxiv. https://doi.org/10.1101/2021.11.09.467869.

Yang, H., Shi, T., Liu, J., Guo, L., Zhang, X., and Yang, S. (2010). A mutant CHS3 protein with TIR-NB-LRR-LIM domains modulates growth, cell death and freezing tolerance in a temperature-dependent manner in Arabidopsis. Plant J. 63, 283–296.

Zdrzałek, R., Kamoun, S., Terauchi, R., Saitoh, H., Banfield M.-J., Wilson R.-A. (2020). The rice NLR pair Pikp-1/Pikp-2 initiates cell death through receptor cooperation rather than negative regulation. PLoS One 15, e0238616.

Zhou, J.-M., Zhang, Y. (2020). Plant immunity: danger perception and signaling. Cell 181, 978–989.

